# scRep: A Latent-Space Self-Distilled Foundation Model for Single-Cell Representation Learning

**DOI:** 10.64898/2026.08.31.747784

**Authors:** Shengjie Wang, Zongyong Hu, Yunlong Bie, Qijin Yin, Hechang Chen, Qiuyi Li

## Abstract

Single-cell foundation models have shown strong potential for learning transferable representations from large-scale transcriptomic data. However, many existing approaches rely on reconstructing masked gene expression values, creating a potential mismatch between observation-space reconstruction and the goal of learning stable biological representations. This challenge is particularly relevant to single-cell RNA sequencing, where sparsity, incomplete gene detection, and technical variation can obscure the underlying biological state. Here, we introduce scRep, a compact latent-space self-distillation framework for single-cell representation learning. Rather than reconstructing raw expression values, scRep aligns differently perturbed views of the same cell through a momentum-updated teacher–student architecture, with self-distillation objectives at both the cell and gene levels. This representation-centered formulation encourages the model to capture biological information that remains stable across incomplete and perturbed transcriptomic observations. Using frozen representations without task-specific fine-tuning, scRep pretrained on approximately 2.8 million cells achieves the strongest overall performance across the evaluated frozen-representation benchmarks, demonstrating strong sample efficiency. A larger-scale scRep model pretrained on 30.72 million cells further demonstrates that the framework remains effective when scaled to a substantially larger and more diverse corpus. Beyond cell identity, scRep prioritizes established marker genes, recovers transcription factor–associated gene programs with cell-type-specific activity, and preserves continuous developmental structure that supports graph-based pseudotime inference. We further show that pretraining performance is closely associated with biological diversity: reducing redundant cells while improving cell-type coverage can match or exceed the performance of larger, less balanced training corpora. Together, these results establish latent-space self-distillation as an effective alternative to expression reconstruction for single-cell foundation modeling and suggest that efficient scaling depends not only on the number of cells, but also on the learning objective and the biological diversity of the pretraining corpus.

## 1 Introduction

High-quality cellular representations are central to single-cell transcriptomic analysis, providing a common basis for tasks such as cell-type annotation, clustering, perturbation analysis, and characterization of cellular states [22, 1, 23, 33, 42, 45, 38]. Rapid advances in single-cell sequencing have generated datasets containing hundreds of millions of cells [30], creating new opportunities to learn transferable representations across tissues, diseases, and experimental contexts. Inspired by foundation models in natural language processing [11, 31, 3], a growing family of single-cell foundation models, including scBERT [44], Geneformer [39], scGPT [8], scFoundation [15], Stack [12], and STATE [1], has sought to leverage these large-scale transcriptomic resources to learn generalizable representations of genes and cellular states.

Many existing single-cell foundation models rely heavily on masked expression reconstruction [44, 39, 15, 8], in which a subset of gene expression values is masked and the model is trained to recover the original observations. Although intuitive, this objective can introduce a mismatch between the pretraining target and the ultimate goal of representation learning. Accurately reconstructing measured expression values does not necessarily require learning the stable biological programs that define cellular identity and state. This distinction is particularly relevant to single-cell RNA sequencing, where observations are high-dimensional, sparse, and incomplete, and are influenced by stochastic transcript capture, sequencing depth, batch effects, donor-specific variation, and experimental protocols [35, 43, 16, 24]. Reconstruction-based objectives may therefore devote substantial modeling capacity to observation-specific variation that is useful for recovering measured expression but not necessarily required for learning transferable representations of the underlying biological state.

In parallel, single-cell foundation models have grown rapidly in pretraining scale. Early models such as scBERT were pretrained on several million cells [44], subsequent models such as scGPT and scFoundation expanded to tens of millions [8, 15], and more recent models including STATE and Stack have scaled beyond one hundred million cells [1, 12]. This trend has substantially advanced the field, but raw cell count alone does not fully characterize the biological information available during pretraining [46, 19, 18]. Large repositories can contain substantial redundancy, with common cell types repeatedly represented while rarer tissues, cellular states, disease contexts, and perturbation conditions remain underrepresented. Consequently, increasing cell count may provide diminishing additional biological information when newly added observations largely repeat states already well represented in the corpus. Effective pretraining scale may therefore depend not only on the number of cells, but also on the biological diversity they contain and on how effectively each cell contributes to representation learning.

These observations motivate an alternative formulation of single-cell pretraining. Multi-view self-supervised learning in computer vision [4, 13, 6, 29] learns representations by enforcing consistency across different transformations or partial views of the same underlying input rather than reconstructing every observed detail. We hypothesized that this principle is particularly well suited to single-cell transcriptomics. A cellular expression profile can be represented as a set of gene-specific observations, with gene identity embeddings specifying which genes are present and expression embeddings describing their measured states [39, 8]. Different stochastic subsets of genes from the same cell then constitute incomplete observations of a shared underlying biological state.

This view is closely aligned with the measurement process of single-cell RNA sequencing. Limited capture efficiency, stochastic transcript detection, sequencing depth, and technology-dependent gene coverage naturally produce partial and technically perturbed observations of the transcriptome. Rather than treating this incompleteness solely as noise to be reconstructed away, it can instead be used as a source of self-supervision. Enforcing consistency across complementary partial views encourages a model to identify biological features that remain stable despite missing genes and variation in measured expression. Stochastic view generation can also expose each biological cell to many distinct observation patterns, increasing the diversity of the training signal without requiring a proportional increase in the number of collected cells [6]. From this perspective, effective pretraining depends jointly on the biological diversity represented across cells and on the diversity of transcriptomic observations generated from each cell.

We therefore introduce scRep, a latent-space self-distillation foundation model for single-cell transcriptomics. scRep formulates pretraining as representation alignment across complementary stochastic views of the same cell rather than reconstruction of raw expression values. During pretraining, gene subsampling, gene dropout, and expression masking generate incomplete transcriptomic views that retain gene identity and observed expression information. A student network learns from latent targets produced by a momentum-updated teacher operating on a more informative view [13, 4], with self-distillation performed at both the cell and gene levels. By optimizing consistency in latent space, scRep turns transcriptomic incompleteness from a quantity to be reconstructed into an invariance signal from which biologically stable representations can be learned.

We evaluate two instances of the same 99.8-million-parameter scRep architecture at different pretraining scales. The primary model, scRep-3M, is pretrained on 2.804 million cells and is used to assess the sample efficiency and biological quality of the learned representations. We additionally train a larger-scale model, scRep-30M, on 30.72 million cells to examine whether the same framework continues to benefit from increased pretraining scale and biological diversity. Using frozen representations without task-specific fine-tuning, scRep-3M achieves the strongest overall performance across the evaluated frozen-representation benchmarks despite its comparatively small pretraining corpus. The larger-scale model further demonstrates that the framework continues to benefit from additional diverse pretraining data.

Overall, scRep advances single-cell foundation modeling in three respects. First, it introduces multi-view latent-space self-distillation as a representation-centered alternative to observation-space reconstruction, with a training formulation naturally matched to the incomplete and technically variable nature of single-cell transcriptomic measurements. Second, it jointly learns transferable cell- and gene-level representations that support downstream analyses without task-specific fine-tuning, including cell clustering and annotation, marker-gene prioritization, transcription factor–associated gene programs, and developmental trajectory analysis. Third, its scaling experiments show that strong representations can be learned from comparatively few cells and that the benefit of additional pretraining data depends strongly on biological diversity rather than raw cell count alone. Together, these results suggest that effective scaling of single-cell foundation models depends not only on how much data are available, but also on the learning objective, how each cell is used during pretraining, and how effectively the training corpus captures biological diversity.

## 2 Results

### 2.1 The scRep self-distillation framework

We developed scRep, a self-distilled Transformer-based foundation model for learning transferable representations from single-cell transcriptomic profiles (Fig. 1). In contrast to approaches that primarily reconstruct masked expression values in the observation space, scRep learns directly in latent space by enforcing consistency across complementary views of the same cell. A student encoder is trained to match latent targets generated by a slowly updated teacher encoder [13, 4]. Through stochastic gene subsampling, gene dropout, expression masking, and view-specific expression binning, the model is exposed to multiple incomplete observations of the same underlying cellular state and is encouraged to retain biological information that remains stable across these perturbations.

**Figure 1:**
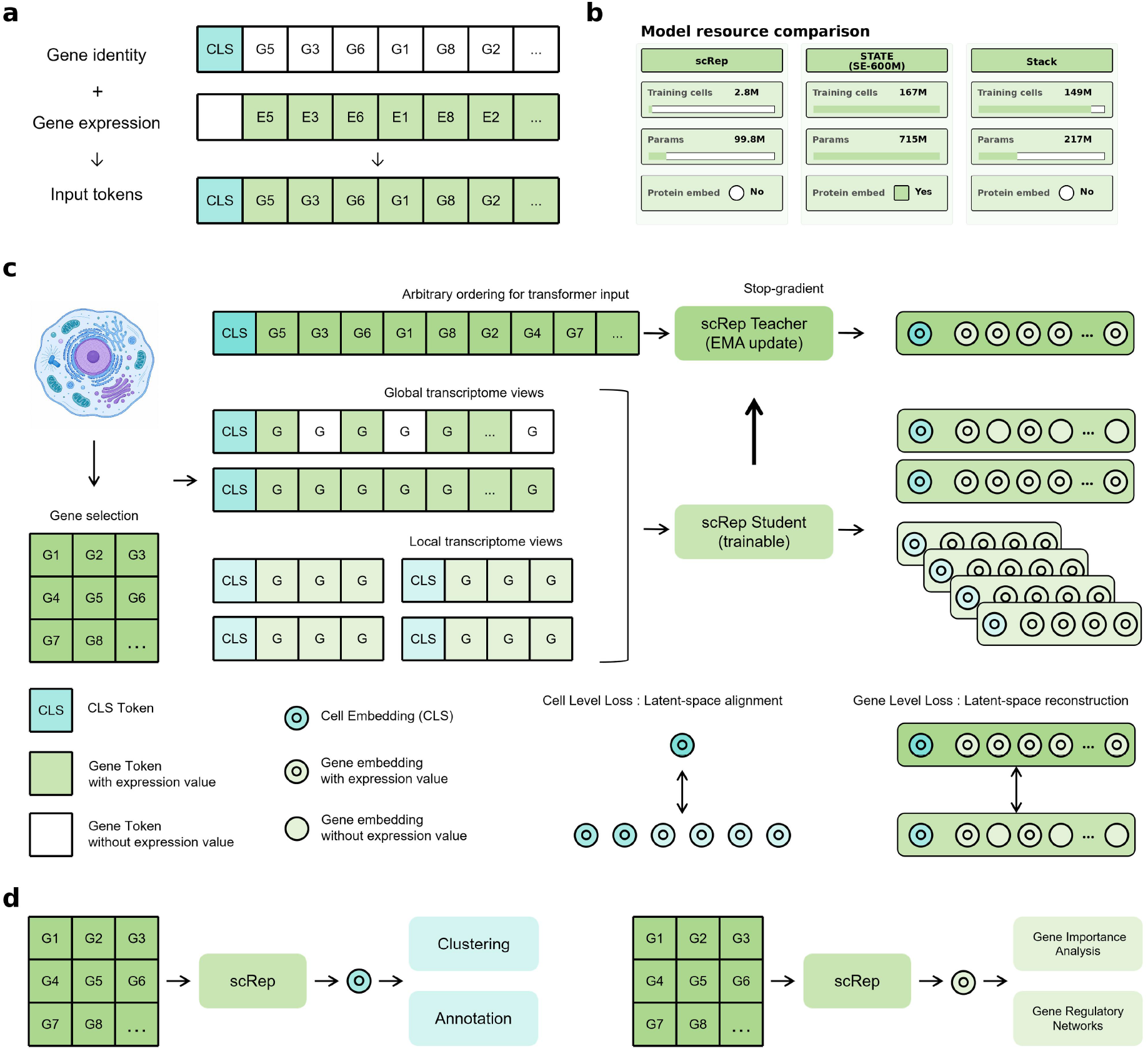
Overview of the scRep framework. a, Expression-aware gene tokenization. Gene identity embeddings are combined with expression embeddings to construct input representations that preserve both gene-specific and cell-state-dependent information. b, Resource comparison of scRep with existing single-cell foundation models. scRep uses substantially fewer parameters and training cells than the models shown. c, Dual-level self-distillation framework. A selected set of genes is arranged as an input sequence with expression-aware embeddings. The teacher receives an unperturbed high-information transcriptome view, whereas the student receives perturbed global and local transcriptome views. Cell-level and gene-level latent-space objectives jointly learn transferable cell and gene representations. d, Biological applications of learned representations. Cell embeddings support unsupervised clustering and cell-type annotation, whereas gene embeddings enable gene importance analysis and embedding-derived regulatory program inference.

scRep represents each cell as a sparse set of observed genes rather than a dense genome-wide expression vector. Each gene token combines a gene identity embedding with an expression-bin embedding, thereby retaining both gene-specific and cell-state-dependent information. Expression values are discretized independently within each augmented view, so that the expression representation reflects the relative expression structure available in that view. A dedicated [CLS] token is prepended to each input and serves as the global cell representation. The resulting tokens are processed by a stack of Transformer encoder blocks [41]; the final [CLS] state provides the cell-level representation, whereas the contextualized gene-token states provide gene-level representations.

During pretraining, scRep constructs asymmetric teacher and student views for each cell. The teacher receives a high-information global view containing up to 2,048 highly expressed observed genes. The student receives six independently perturbed views comprising two global views and four local views. Student global views are generated from the teacher gene pool through stochastic gene subsampling and dropout, whereas local views contain fewer genes and undergo stronger subsampling. These asymmetric views provide observations of the same biological cell at different levels of completeness, requiring the student to recover information that is consistent across partial transcriptomic measurements.

Student global views are additionally used for expression-masking-based gene representation learning. For selected global views, a subset of genes has its expression information masked before expression-bin construction. Masked genes retain their gene identities but are assigned a dedicated mask-bin identifier, while expression bins for the remaining genes are calculated using only unmasked expression values. This prevents masked expression values from leaking into the input through view-level bin assignments and requires the student to infer their contextual representations from gene identity and the surrounding transcriptomic context.

scRep optimizes complementary self-distillation objectives at the cell and gene levels. At the cell level, the [CLS] representation from the teacher view and those from each of the six student views are passed through cell-level projection heads. Rather than directly matching hidden embeddings, scRep aligns the resulting teacher and student output distributions. Teacher predictions are centered and temperature-sharpened to provide stable targets [4], and the student is optimized against these targets using cross-entropy. The teacher receives no gradient updates and is instead updated as an exponential moving average of the student network [13, 4]. The cell-level objective therefore encourages the global cell representation to remain consistent across substantially different partial observations of the same cell.

In parallel, gene-level self-distillation directly supervises contextualized gene-token representations. For masked genes that occur in both the teacher and student views, teacher and student token states are matched by gene identity rather than sequence position. This is important because genes are independently sampled and shuffled across views. The student representation of each masked gene is then trained to match the corresponding teacher target through a separate gene-level projection head. In this way, the gene-level objective encourages gene representations to encode their cellular context rather than only static gene identity, while remaining robust to missing or perturbed expression information.

The two objectives are optimized jointly:

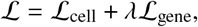

where ℒ_cell_ denotes cell-level self-distillation based on [CLS] representations and ℒ_gene_ denotes token-level self-distillation for matched genes. We set λ = 1, assigning equal weight to the two objectives. Together, these objectives encourage scRep to preserve both global cellular state and contextual gene-level structure without explicitly reconstructing the original expression profile.

After pretraining, the momentum teacher encoder is used as a frozen feature extractor for downstream analyses, with no task-specific fine-tuning of the pretrained encoder. We trained two instances of the same 99.8-million-parameter architecture at different pretraining scales. scRep-3M, pretrained on 2.804 million cells, serves as the primary model for evaluating representation quality and sample efficiency. We additionally trained scRep-30M on 30.72 million cells as a larger-scale extension to examine whether the same framework continues to benefit from increased pretraining scale and biological diversity.

### 2.2 scRep learns biologically meaningful cell representations

A central goal of single-cell foundation models is to learn representations that preserve cellular identity while remaining transferable across diverse biological contexts. We therefore evaluated frozen scRep cell embeddings in two complementary downstream settings: unsupervised clustering of cellular populations and reference-based cell-type annotation [40, 26, 20]. Together, these analyses assess whether the learned representation space captures biologically meaningful cellular structure and supports label transfer without task-specific fine-tuning.

### 2.2.1 Biological structure emerges in the learned latent space

We first assessed whether frozen scRep embeddings preserve cellular organization without using cell-type labels to train the representation. We evaluated six independent benchmark datasets spanning diverse tissues and disease contexts: Glioblastoma, Brain, Kidney, Eye, Pancreas, and Heart. For each dataset, we performed Leiden clustering on the frozen cell embeddings and compared the resulting clusters with curated cell-type annotations. For evaluation, Leiden clustering was calibrated to match the number of annotated cell types.

Across the six datasets, scRep achieved the highest mean clustering agreement among the evaluated models, with a mean normalized mutual information (NMI) of 0.859 and adjusted Rand index (ARI) of 0.821 (Fig. 2a). Strong performance was observed in Glioblastoma (NMI = 0.944; ARI = 0.957; Fig. 2b) and Eye (NMI = 0.918; ARI = 0.879; Fig. 2c), indicating that scRep resolves cellular heterogeneity across distinct tissue and disease settings.

**Figure 2:**
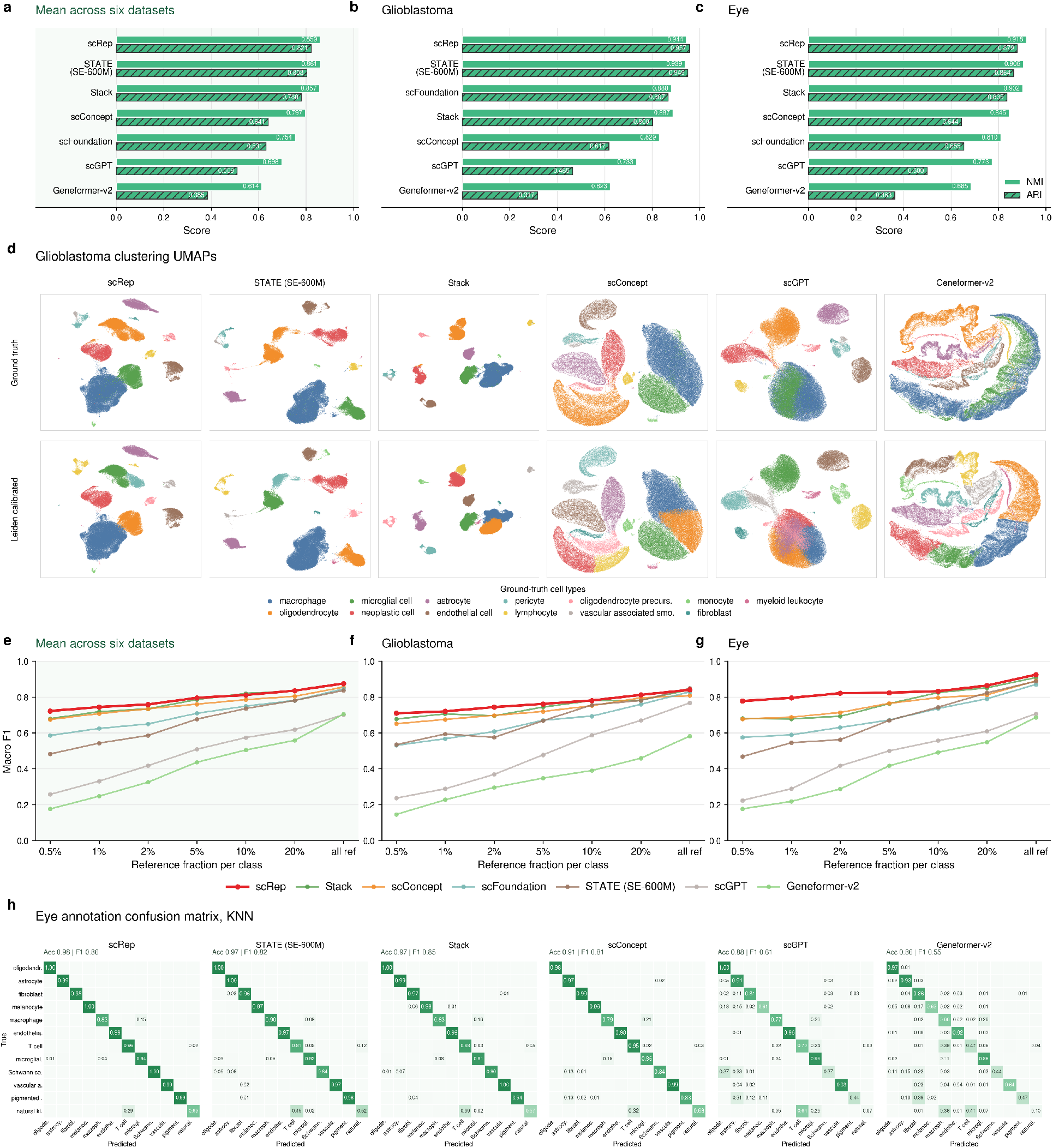
Benchmarking biological representation quality of scRep. a-c, Ranking of unsupervised clustering performance measured by normalized mutual information (NMI) and adjusted Rand index (ARI). Panel a summarizes the mean performance across six benchmark datasets, whereas panels b and c show results for the Glioblastoma and Eye datasets, respectively. d, UMAP visualizations of clustering results on the Glioblastoma dataset for scRep, STATE, Stack, scConcept, scGPT, and Geneformer-v2. For each model, ground-truth cell-type annotations are shown in the upper row and Leiden-calibrated clustering assignments in the lower row. e–g, KNN cell-type annotation performance, measured by macro-F1, as a function of the reference fraction per cell type. Panel e summarizes the mean performance across six benchmark datasets, whereas panels f and g show results for the Glioblastoma and Eye datasets, respectively. h, Confusion matrices for KNN cell-type annotation on the Eye dataset, comparing predicted and ground-truth cell types for the same six models.

Consistent with the quantitative results, UMAP visualizations of the Glioblastoma dataset showed that cells with the same annotated identity occupied coherent regions of the scRep latent space, while Leiden-calibrated clusters closely recapitulated the corresponding cell-type structure (Fig. 2d). These results show that scRep embeddings preserve biologically meaningful cellular organization in an unsupervised clustering setting.

#### 2.2.2 Frozen embeddings support accurate cell-type annotation

We next examined whether the biological structure encoded by scRep embeddings supports cell-type annotation across independent biological samples. We used frozen embeddings for KNN-based label transfer, in which annotated reference cells were used to infer the identities of held-out query cells. This evaluation does not involve task-specific fine-tuning of scRep.

To mimic realistic annotation settings, reference and query cells were separated by donor or sample identity, such that biological groups were not shared between the two partitions. We then varied the fraction of labeled reference cells available for each cell type. Specifically, each value on the horizontal axis denotes the proportion of reference cells retained per cell type, ranging from 0.5% to 20%; “all ref” denotes the use of all available reference cells in the reference partition.

Across the six datasets, scRep achieved the highest mean macro-F1 score at most reference fractions, including the low-reference settings from 0.5% to 5% and the full-reference setting (Fig. 2e). At the most stringent setting, in which only 0.5% of available reference cells were retained per cell type, scRep achieved a mean macro-F1 score of 0.7222. Depending on the dataset and cross-validation fold, this setting corresponded to approximately 3–20 reference cells per cell type.

With 5% labeled reference cells per cell type, scRep attained a mean accuracy of 0.9001 and a mean macro-F1 score of 0.7955. With the full reference set, performance increased to a mean accuracy of 0.9328 and a mean macro-F1 score of 0.8752. Notably, reducing the reference set from the full-reference setting to 0.5% per cell type decreased the mean macro-F1 of scRep by only 0.1530, representing the smallest performance reduction among the evaluated models. These results indicate that scRep embeddings remain highly informative when labeled reference data are scarce and support robust annotation in low-reference settings. The same overall pattern was observed in both the Glioblastoma and Eye datasets (Fig. 2f,g).

The Eye-dataset confusion matrices further showed that scRep accurately distinguished most cell types and exhibited limited off-diagonal confusion relative to several baseline models (Fig. 2h). Collectively, these results demonstrate that frozen scRep embeddings contain transferable cell-identity information and support accurate reference-based annotation even when labeled reference data are limited.

### 2.3 Pre-training data diversity is associated with representation quality beyond data scale

We next asked whether improvements in downstream representation quality could be explained by pre-training corpus size alone [10], or whether the biological diversity of the pre-training data provided additional explanatory value. We quantified the cell-type-aware diversity of each pre-training corpus using a diversity score that assigns diminishing marginal weight to increasingly abundant cell populations (Methods), and compared this quantity with downstream clustering performance across six independent benchmark datasets.

Across the primary pre-training scaling series, increases in corpus size were accompanied by increases in both biological diversity and downstream representation quality (Fig. 3a–c). Tabula Sapiens, containing 1.136 million cells, achieved a mean clustering performance of 0.76099. Expanding the corpus to 2.000 million cells using additional cellxgene data increased performance to 0.82493, while the 2.804-million-cell corpus further improved performance to 0.83161. These gains were accompanied by increases in cell-type-aware diversity from 1290.94 to 5917.39 and 6400.35, respectively.

**Figure 3:**
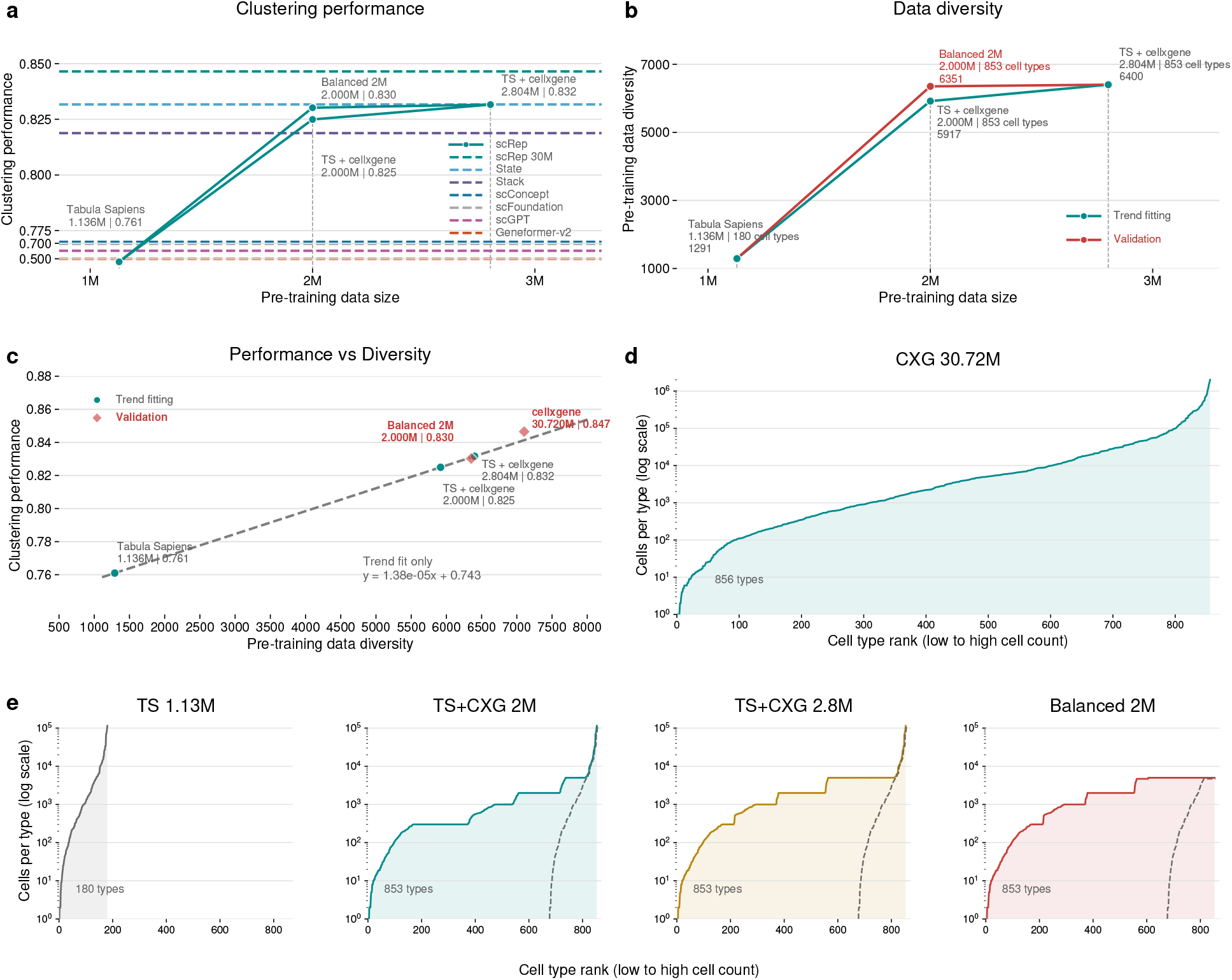
Effects of pre-training data scale and diversity on scRep performance. a, Clustering performance of scRep models pre-trained on different data compositions, measured as mean calibrated ARI and NMI across downstream clustering benchmarks. Dashed horizontal lines indicate the performance of scRep pre-trained on 30.72M cellxgene cells and representative baseline models. b, Cell-type-aware diversity scores of pre-training datasets across increasing data scales. Higher values indicate broader cell-type coverage and greater representation of underrepresented populations. c, Relationship between pre-training data diversity and clustering performance. The fitted trend is estimated from the main scaling series, with the Balanced 2M dataset shown separately as a held-out validation case. d, Cell-type abundance distribution of the 30.72M profiles used to train scRep-30M. Cell types are ranked by profile count to illustrate the long-tailed distribution of the training composition. e, Cell-type abundance distributions of smaller pre-training datasets, including Tabula Sapiens, TS + cellxgene 2M, TS + cellxgene 2.8M, and the Balanced 2M dataset. Dashed curves indicate the corresponding Tabula Sapiens-derived distributions where applicable.

Importantly, matched-scale comparisons showed that cell number alone did not fully account for the observed differences. The coverage-balanced 2M corpus (Balanced 2M), constructed by reducing redundant cells while retaining broader and more balanced cell-type coverage, contained approximately the same number of cells as the direct 2M corpus but had substantially higher diversity (6351.41 versus 5917.39). This increase in diversity was accompanied by improved clustering performance (0.83020 versus 0.82493; Fig. 3a–c).

A complementary comparison was observed between the Balanced 2M and the original 2.804M corpus. Despite containing approximately 0.8 million fewer cells, the Balanced 2M corpus had a diversity score close to that of the larger corpus (6351.41 versus 6400.35) and achieved nearly identical downstream performance (0.83020 versus 0.83161). Thus, removing redundant observations while preserving biological coverage retained most of the representation quality obtained from the larger training corpus.

The relationship between biological diversity and representation quality was further supported by the scaling trend across the primary pre-training series (Fig. 3c). The Balanced 2M corpus, which was not used to fit this trend, closely followed the relationship predicted from the primary scaling datasets, deviating by only −0.00073 in clustering performance. The substantially larger 30.72-million-cell corpus showed the highest diversity score (7103.87) and the strongest observed clustering performance (0.84650), while remaining broadly consistent with the same association between greater pre-training diversity and stronger downstream representations. Because this large-scale run differed from the primary scaling series in both data composition and training configuration, we interpret it as supporting evidence for the broader trend rather than as a controlled extension of the smaller-scale comparison.

Together, these results indicate that the effective scale of single-cell pre-training cannot be characterized by cell number alone. Increasing the number of cells was most beneficial when it expanded biological coverage, whereas adding additional observations from already well-represented populations provided smaller gains. The matched-scale comparison between the two 2M corpora, together with the near-equivalent performance of the Balanced 2M and original 2.804M corpora, suggests that biological diversity is an important determinant of representation quality and that reducing redundancy can improve the efficiency of single-cell foundation-model pre-training.

### 2.4 scRep captures biologically meaningful gene importance and regulatory programs

Beyond representing cellular identities, a useful single-cell foundation model should also preserve biologically meaningful information at the gene level. We therefore examined scRep from two complementary perspectives. First, we asked whether genes known to define specific cell identities exerted greater influence on the learned cell representations. Second, we tested whether the contextual organization of scRep gene embeddings captured transcription factor (TF)–associated gene programs with cell-type-specific activity.

#### 2.4.1 Gene importance scores prioritize cell-type marker genes

We first tested whether genes that are biologically important for defining cellular identity also contribute strongly to scRep cell representations. Using a perturbation-based sensitivity score that measures the change in the cell embedding after removing the expression information of an individual gene while retaining its identity (Methods), we evaluated known marker genes across 16 cell types in the Eye dataset [28, 25, 27, 21].

Known marker genes consistently ranked above other genes according to their conditional perturbation effect across cell types (Fig. 4a). This enrichment was also observed after accounting for how frequently each gene was detected within the corresponding cell population (Fig. 4b), indicating that marker prioritization was not restricted to genes exerting large effects in only a small subset of cells.

**Figure 4:**
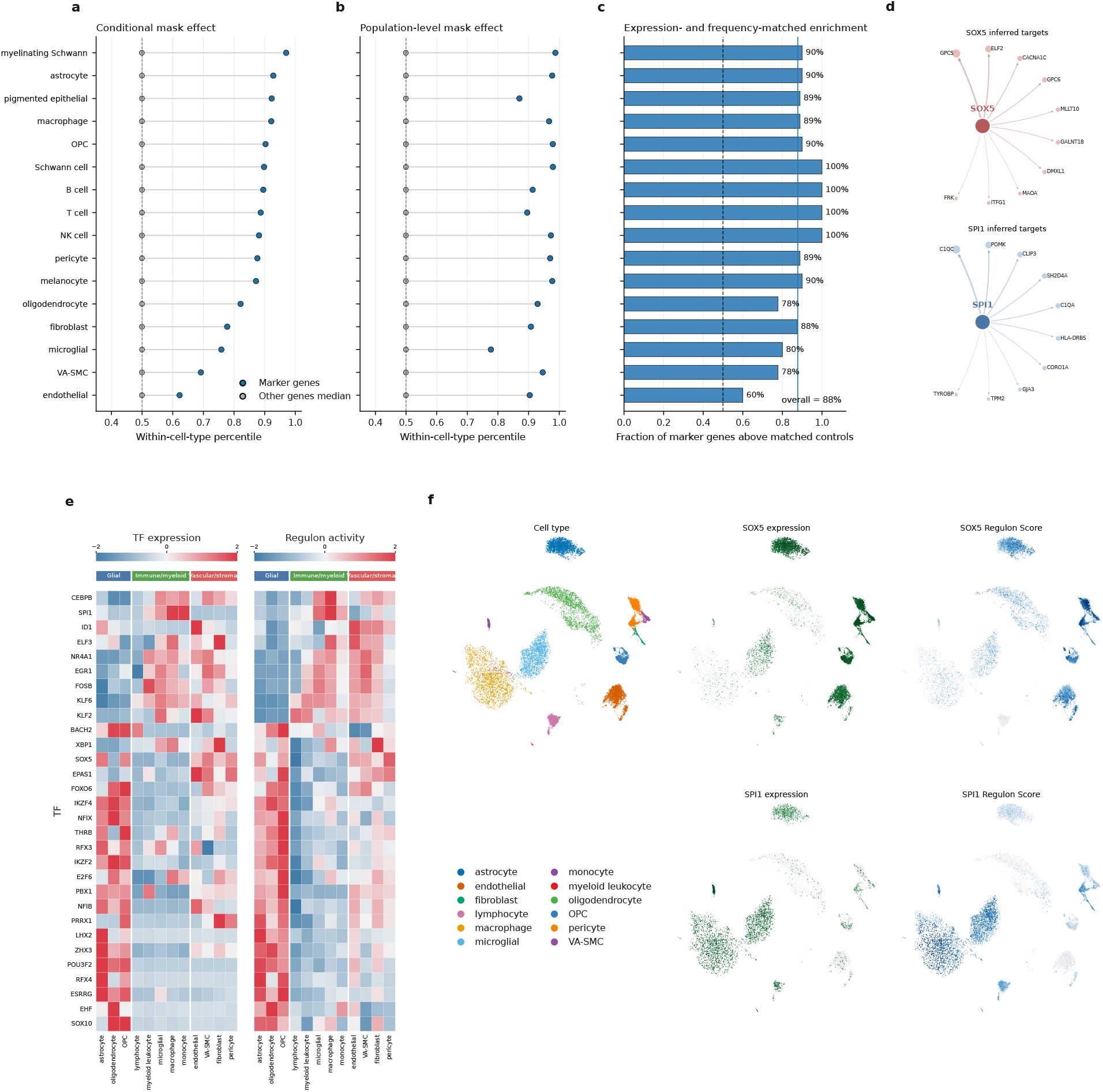
Gene importance and transcription factor regulon characterization. a-c, Evaluation of cell-type marker gene prioritization using scRep-derived gene representations. Panels a and b show within-cell-type percentile ranks of marker genes compared with the median of other genes, measured by conditional mask effect and population-level mask effect, respectively. Panel c shows the fraction of marker genes ranked above expression- and frequency-matched control genes. d, Embedding-derived TF-associated networks centered on two representative transcription factors, SOX5 and SPI1, in the Glioblastoma dataset. For visualization, the 10 highest-scoring inferred target genes are shown for each transcription factor. e, Heatmaps of TF expression and regulon activity across annotated Glioblastoma cell types. Cell types are grouped by broad lineage categories, including glial, immune/myeloid, and vascular/stromal populations. f, UMAP visualizations of the Glioblastoma dataset colored by annotated cell type, SOX5 and SPI1 expression, and SOX5 and SPI1 regulon activity. Together, these panels show that inferred TF programs are spatially structured and cell-type specific. Abbreviations: OPC, oligodendrocyte precursor cell; NK cell, natural killer cell; VA-SMC, vascular-associated smooth muscle cell.

Because marker genes are often highly expressed or broadly detected, we further compared each marker with non-marker genes matched for expression abundance and detection frequency. Across cell types, approximately 88% of marker genes ranked above their matched controls (Fig. 4c). Thus, the preferential sensitivity of scRep representations to established marker genes could not be explained solely by their expression level or prevalence.

Together, these results show that scRep cell representations are selectively sensitive to gene-level signals associated with cellular identity, providing an interpretable readout of which observed gene-expression states most strongly influence the learned representation.

#### 2.4.2 Gene embeddings reveal cell-type-specific regulatory programs

We next asked whether the contextual structure encoded by scRep gene embeddings captured biologically coherent TF-associated gene programs. Using an embedding-similarity-based procedure applied to the Glioblastoma dataset (Methods), we constructed putative TF–target networks from contextualized gene representations, following the general strategy of previous gene-embedding-based regulatory analyses [17].

The resulting networks showed distinct TF-specific neighborhoods (Fig. 4d). For example, the SOX5-associated network included high-similarity candidate targets such as GPC5, GPC6, CACNA1C, ELF2, MLLT10, and GALNT18, whereas the SPI1-associated network included several genes linked to immune and myeloid states, including C1QA, C1QC, CORO1A, TYROBP, and HLA-DRB5. These contrasting neighborhoods indicate that the geometry of scRep gene embeddings organizes transcription factors and candidate target genes into distinct functional programs.

We next examined whether these embedding-derived target sets exhibited cell-type-specific activity. Across 30 selected transcription factors, regulon activity showed pronounced variation across glial, immune/myeloid, and vascular/stromal populations and broadly corresponded to the expression patterns of the associated TFs (Fig. 4e). At the same time, regulon activity was not identical to TF expression, providing a complementary readout of the expression state of the inferred downstream gene program.

Distinct lineage-associated programs were apparent across cell types. TFs associated with glial, immune/myeloid, and vascular/stromal populations showed corresponding cell-type-specific regulon activity patterns, indicating that the inferred target sets were not uniformly active across cellular populations.

At single-cell resolution, SOX5- and SPI1-associated regulon activity occupied strongly contrasting regions of the Glioblastoma embedding space (Fig. 4f). SPI1 regulon activity was concentrated predominantly in immune and myeloid populations, consistent with both SPI1 expression and its established role in myeloid regulatory programs. In contrast, SOX5-associated activity was enriched in a distinct region of the cellular landscape, separating it from the SPI1-associated myeloid state [9, 34, 32, 37, 36].

The broad spatial correspondence between regulon activity and TF expression, together with substantial cell-to-cell variation between the two quantities, suggests that the embedding-derived regulon scores capture coordinated downstream transcriptional states rather than simply reproducing TF transcript abundance.

Collectively, these analyses reveal two complementary forms of gene-level biological information captured by scRep. Perturbation-based sensitivity preferentially highlights established cell-type markers even after controlling for expression abundance and detection frequency, whereas contextualized gene embeddings organize TF-associated gene programs whose activities vary coherently across cellular populations. These findings indicate that scRep retains biologically structured information at both the individual-gene and gene-program levels.

### 2.5 scRep embeddings recover developmental ordering from latent representations

We next asked whether the continuous biological structure encoded by scRep embeddings could support the recovery of developmental progression. We therefore applied graph-based diffusion pseudotime analysis to frozen scRep representations in two human single-cell datasets with independent temporal or state annotations (Methods) [14].

We first examined a human peri-implantation embryo dataset sampled at D6, D8, D10, D12, and D14 (Fig. 5a–c). scRep-3M produced a continuous pseudotime gradient across the cellular embedding (Fig. 5a) that closely followed the independently recorded developmental-day pattern (Fig. 5b). Pseudotime distributions shifted progressively toward larger values from D6 to D14, yielding a cell-level Spearman correlation of *ρ* = 0.794 between inferred pseudotime and developmental day (Fig. 5c).

**Figure 5:**
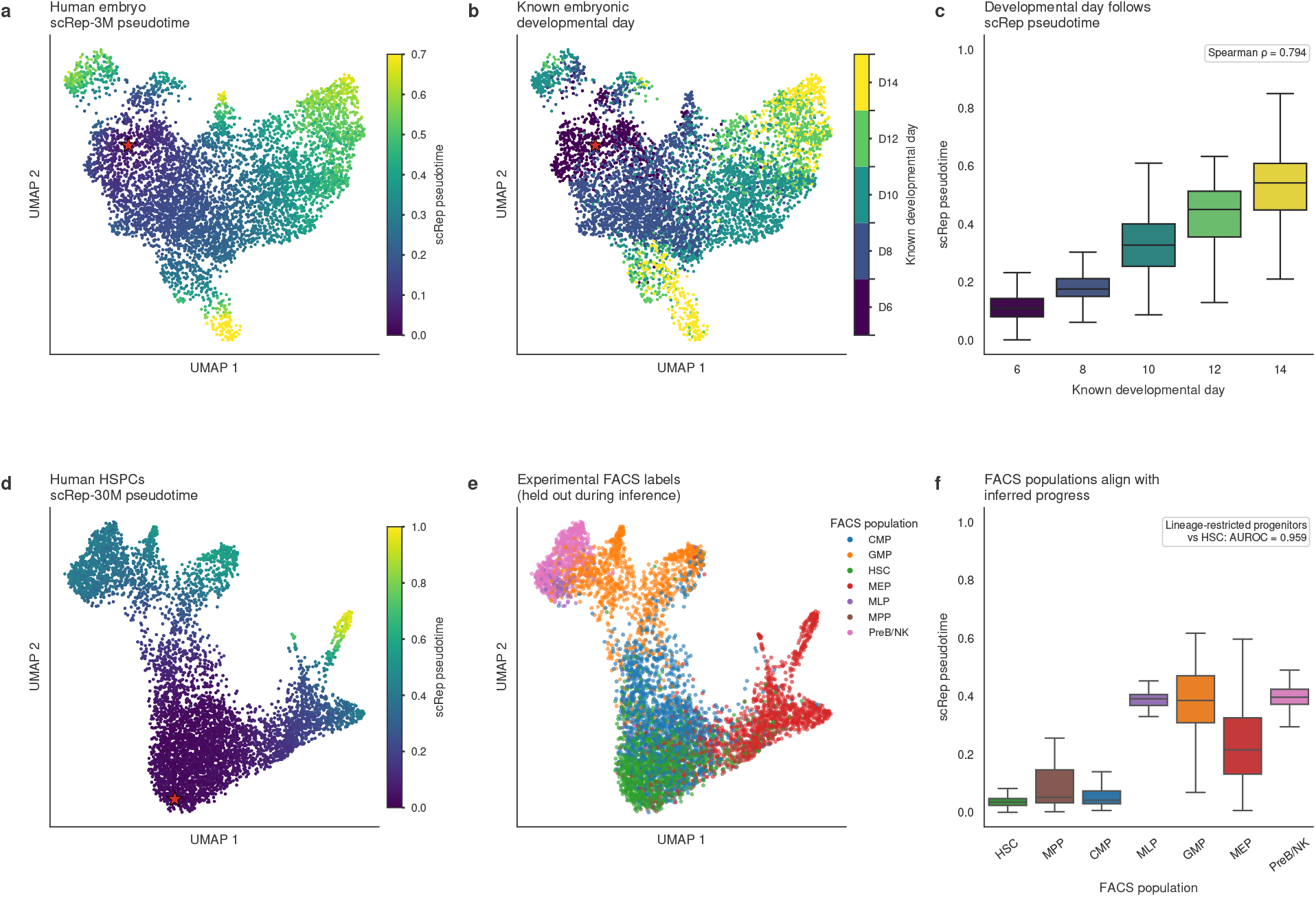
scRep embeddings recover developmental ordering from frozen representations. a, scRep-3M-derived pseudotime in human peri-implantation embryos. b, Known embryonic developmental day projected onto the same embedding; developmental day was withheld during inference. c, Distribution of inferred pseudotime across developmental days. Spearman’s *ρ* between scRep pseudotime and developmental day was 0.794. d, scRep-30M-derived pseudotime for human HSPCs; the star denotes the selected early-state root. e, Experimental FACS populations projected onto the same embedding. FACS labels were withheld during trajectory inference. f, Distribution of inferred pseudotime across FACS populations. Lineage-restricted progenitors (GMP, MEP and PreB/NK) were distinguished from HSCs with an AUROC of 0.959.

Developmental day was withheld from representation learning and trajectory construction, except for selecting an early-state root from D6 cells to orient the inferred trajectory. Thus, the observed agreement reflects recovery of temporal ordering from the geometry of the scRep embedding beyond this minimal directional anchor.

We next evaluated scRep-30M in human haematopoietic stem and progenitor cells (HSPCs), for which experimentally defined FACS populations provide an independent reference for developmental state (Fig. 5d–f). In this dataset, trajectory inference was performed without using FACS population labels, and the root was selected independently using an HSC marker-expression score (Methods).

The resulting pseudotime again formed a continuous gradient across the scRep embedding (Fig. 5d). When the held-out FACS labels were overlaid after inference, HSCs and multipotent progenitors were concentrated at lower pseudotime, whereas MLP, GMP, and PreB/NK populations occupied later regions; MEP cells showed a broader intermediate distribution (Fig. 5e,f). Consistent with this shift, inferred pseudotime distinguished lineage-restricted progenitors (GMP, MEP, and PreB/NK) from HSCs with an AUROC of 0.959 (Fig. 5f). This AUROC summarizes the ordering induced by the inferred pseudotime and does not result from a supervised classifier trained on FACS labels.

The HSPC populations did not exhibit a single strict linear ordering, consistent with the branched structure of haematopoietic differentiation and the fact that FACS populations represent experimentally enriched states rather than consecutive positions along a single lineage. Together, the embryo and HSPC analyses show that scRep embeddings preserve continuous developmental structure: inferred trajectories recapitulate known temporal progression after minimal root anchoring in embryos and separate independently annotated early and lineage-restricted haematopoietic states without using FACS labels during trajectory inference.

## 3 Discussion

In this study, we introduce scRep, a latent-space self-distillation framework for learning transferable representations from single-cell transcriptomes. Rather than reconstructing masked expression values, scRep aligns representations from complementary stochastic views of the same cell through a momentum-updated teacher–student architecture. The resulting frozen embeddings support unsupervised clustering and cell type annotation across diverse datasets, while also preserving biologically meaningful gene-level information and continuous developmental structure. These results suggest that representation-centered self-supervision provides an effective alternative to observation-space reconstruction for single-cell foundation modeling.

Our scaling experiments further indicate that pretraining performance depends on biological diversity rather than cell number alone [10]. Increasing dataset size improved representation quality when it expanded cell-type coverage, whereas adding redundant cells produced smaller gains. Consistently, a deduplicated 2M-cell dataset matched or exceeded the performance of a larger non-deduplicated corpus, and its observed performance was consistent with the trend captured by our cell-type-aware diversity score. These findings suggest that future single-cell foundation models may benefit not only from larger datasets, but also from more deliberate corpus design that improves biological coverage and reduces redundancy.

Beyond cell-level representation, scRep also captures biologically structured gene information. Gene perturbation analysis preferentially assigned high importance to established marker genes even after controlling for expression abundance and detection frequency, while contextualized gene embeddings supported the inference of TF-associated gene programs with cell-type-specific regulon activity. In addition, graph-based pseudotime inferred from scRep embeddings recovered developmental progression in human embryos and separated early from lineage-restricted states in human haematopoietic progenitors. Together, these results indicate that the learned representation space retains both discrete cellular identities and continuous biological variation, while also encoding functional relationships at the gene level.

Several limitations remain. The current benchmarks cover only a subset of tissues, technologies, species, and perturbational settings, and broader evaluation will be required to define the limits of zero-shot transfer. Our diversity score is also intentionally simple and depends on cell-type annotations, which do not capture all dimensions of biological variation. In addition, embedding-derived TF–target relationships should be interpreted as putative functional associations rather than causal regulatory interactions and will require validation with orthogonal regulatory or perturbation data. Despite these limitations, our results support two complementary principles for efficient single-cell foundation modeling: learning objectives should be aligned with robust biological representation rather than noisy observation reconstruction, and pretraining scale should prioritize biological diversity rather than cell count alone.

## 4 Methods

### 4.1 Input tokens

For each cell, scRep represents the transcriptome as a variable-length sequence of detected genes rather than a fixed ordered gene-expression vector. Let 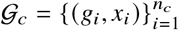 denote the non-zero entries of cell *c*, where *g*_*i*_ is a gene identity and *x*_*i*_ is its expression value. Zero-expression genes are not included in the encoder input sequence. A special [CLS] token is prepended to every sequence, and sequences within a mini-batch are right-padded to the maximum sequence length of that batch. Padding positions are excluded from self-attention.

The model uses a global vocabulary of 19,238 biological genes collected from the training corpus, together with two special gene tokens, yielding a gene vocabulary size of 19,240. Input genes are randomly permuted after view construction. Since no positional embedding is used, the encoder operates on a set of expressed genes rather than on an arbitrary gene order.

#### Gene tokens

A global gene vocabulary was constructed by scanning the gene names in all training h5ad files. Each unique gene identifier was assigned one integer token ID shared across datasets. The gene vocabulary reserves ID 0 for [PAD] and ID 1 for [CLS], while biological genes are assigned IDs from 2 onward.

For a cell *c*, only genes with non-zero expression are retained. The resulting gene-ID sequence is

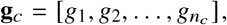

and the encoder input is

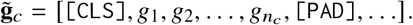

The final representation of the [CLS] token is used as the cell-level embedding.

During self-supervised pre-training, we first retained at most the 2,048 highest-expression genes in each cell as the teacher gene pool. The teacher received one global view containing all genes in this pool. The student received two global views and four local views sampled from the same teacher gene pool. Each global student view was constructed by uniformly sampling 60–85% of the available teacher-pool genes, subject to a target minimum of 1,000 genes when sufficient genes were available. Each local view contained 256–768 sampled genes. Independent gene dropout was subsequently applied with probability 0.10 to student global views and 0.20 to student local views, while preserving the corresponding minimum numbers of genes. Genes were shuffled independently in every view.

#### Expression values and discretization

The training data use the expression matrix stored in adata.X, which was treated as pre-normalized log-transformed expression data. Expression discretization was performed independently for each cell and each augmented view. Therefore, an expression bin indicates the relative expression level of a gene within the current view, rather than a fixed absolute expression interval shared across cells.

For an unmasked view containing *m* genes with expression values 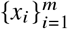, we partitioned the observed expression values into up to *B* = 50 approximately equal-frequency bins. Expression^*i*=^v^1^alues were ordered from low to high. The binning procedure adaptively merged adjacent expression levels or split tied-value groups when necessary, such that bins contained approximately equal numbers of genes. Each positive-expression gene was assigned an expression token

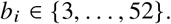

The expression-token vocabulary contained 53 entries: [PAD] (0), [MASK] (1), a reserved zero-expression token (2), and 50 positive-expression bins (3–52). As only detected genes were included in encoder inputs, the zero-expression token was not normally used in input views.

We used DINOv2-style stochastic masking on student global views [29]. Across all global student views in each distributed mini-batch, 50% of views were selected for expression masking. For every selected view, the masking ratio was independently sampled as

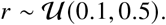

Each gene in the selected view was independently masked with probability *r*, and its expression-bin token was replaced with the [MASK] token. The gene-identity token remained visible. Thus, the model was required to learn cellular context and expression-aware gene relationships from the visible genes and unmasked expression values.

For masked global student views, masking was performed before binning. Specifically, expression bins were computed using only the unmasked genes in the selected view, while masked genes were directly assigned the [MASK] expression token. This design prevents the expression values of masked genes from influencing the bin boundaries of visible genes. Global views not selected for masking, all four local student views, and the teacher global view used unmasked expression-bin tokens.

#### Token embeddings

Each input position was represented by the sum of a learned gene-identity embedding and a learned expression-token embedding. Let

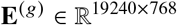

denote the gene embedding table and

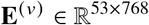

denote the expression-token embedding table. For biological gene *g*_*i*_ with expression token *b*_*i*_, its input representation was

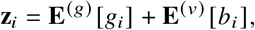

where **z**_*i*_ ∈ ℝ^768^.

For the prepended [CLS] token, the gene ID was 1 and the expression ID was [PAD] (0), giving

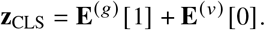

Because the expression-padding embedding was fixed to zero by the embedding layer’s padding mechanism, the initial [CLS] representation was determined solely by its learned gene-token embedding. Similarly, padding positions used gene and expression ID 0 and were excluded from attention computation.

No positional embeddings were introduced. Together with random gene shuffling, this ensured that the input representation and Transformer encoder were insensitive to the arbitrary ordering of genes within each cell view.

### 4.2 Cell and gene representation modeling

#### scRep transformer encoder

scRep employs an encoder-only Transformer to model dependencies among genes expressed in the same cell. For an input sequence of *L* tokens, the encoder maps the input embeddings

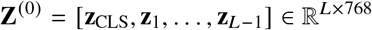

to contextualized hidden states

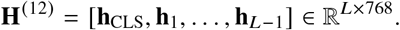

The encoder contains 12 Transformer blocks, with hidden dimension *d* = 768 and 12 attention heads. Thus, each attention head has dimension 64. In each block, multi-head self-attention is followed by a feed-forward network (FFN) with an intermediate dimension of 4*d* = 3,072. The FFN consists of a linear projection from 768 to 3,072 dimensions, a Gaussian error linear unit (GELU) activation, and a linear projection back to 768 dimensions. All linear projections in the Transformer backbone are bias-free.

Each Transformer block uses pre-normalization with root-mean-square normalization (RMSNorm). Given the input **H**^(ℓ−1)^ to layer ℓ, the layer is defined as

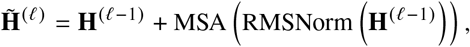

followed by

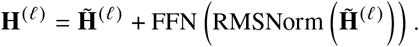

A final RMSNorm layer is applied after the 12th Transformer block. Residual connections are used around both the self-attention and FFN sublayers. The dropout probability is set to 0.0 throughout the encoder.

Unlike language models, scRep does not use positional embeddings. Gene order within a cell has no biological meaning, and input genes are randomly shuffled during view construction. Instead, each gene is identified by its learned gene embedding, which provides the gene-specific identity information required to distinguish tokens without imposing an artificial sequential order. Consequently, the encoder is insensitive to the arbitrary ordering of genes in the input sequence.

Within each Transformer layer, self-attention enables every gene token to attend to all non-padding gene tokens and to the [CLS] token in the same cell. This mechanism allows the representation of a gene to be updated according to its co-expressed genes, their relative expression levels, and the global transcriptional context of the cell. Therefore, the encoder models genes as context-dependent components of a cellular gene-expression program rather than as independent features.

#### Cell representation

For a cell *c*, the cell representation is defined as the final hidden state of the prepended [CLS] token:

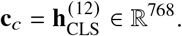

Because the [CLS] token participates in self-attention with all expressed-gene tokens, its final hidden state aggregates information from the complete input gene set and their discretized expression values.

During pre-training, the [CLS] hidden state from each teacher or student view is passed to a cell-level projection head. The projection head maps the 768-dimensional backbone representation to the space used for cell-level self-distillation. This projection head is used only to define the pre-training objective and is not used for downstream analyses.

For downstream tasks, we use the 768-dimensional [CLS] hidden state directly from the teacher encoder backbone as the cell embedding. The teacher encoder is selected by default at inference because it is the exponentially moving averaged version of the student encoder. The downstream cell embedding is taken before the cell-level projection head and is not explicitly ℓ_2_-normalized by the backbone. Although the final RMSNorm controls the scale of hidden states, it does not impose unit Euclidean norm; task-specific downstream analyses may optionally apply normalization when required.

#### Contextualized gene representation

For gene *g*_*i*_ in cell *c*, scRep defines its gene representation as the final hidden state at the corresponding gene-token position:

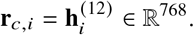

This representation is distinct from the initial learned gene embedding **E**^(*g*)^ *g*_*i*_. The initial embedding encodes the global identity of a gene, whereas **r**_*c,i*_ is a contextualized representation that depends on the complete input cell:

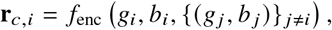

where *b*_*i*_ denotes the expression token of gene *g*_*i*_.

Thus, the same gene can obtain different representations across cells according to its expression level, co-expressed genes, cell state, and cell type. For example, a gene expressed in immune cells and the same gene expressed in epithelial cells may have distinct contextualized representations even though both originate from the same learned gene-identity embedding.

During pre-training, contextualized gene-token states can additionally be passed through a gene-level projection head for token-level representation learning. This projection head is used only for the pre-training objective; the contextualized 768-dimensional backbone token state is the gene representation used for downstream gene-level analyses.

### 4.3 Latent-space self-distillation pretraining

#### Asymmetric teacher–student framework

scRep is pretrained using an asymmetric teacher–student self-distillation framework designed to learn stable cellular and gene representations from incomplete and perturbed transcriptomic observations. The student and teacher networks share the same architecture, consisting of an encoder-only Transformer backbone, a cell-level projection head, and a gene-level projection head, but differ in their optimization roles and input views.

Let ***θ*** and ***ξ*** denote the parameters of the student and teacher networks, respectively. The student is optimized by back-propagation, whereas the teacher receives no gradients. Instead, the teacher parameters are updated after each optimization step as an exponential moving average (EMA) of the student parameters:

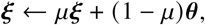

where *μ* is the teacher momentum coefficient. The teacher therefore acts as a slowly evolving target network that aggregates historical student states and provides stable targets for self-distillation.

The framework is also asymmetric in the information presented to the two networks. For each cell, the teacher receives one relatively complete global transcriptomic view, whereas the student receives six independently augmented views comprising two global views and four local views. Student views contain fewer genes and undergo stronger perturbations, including stochastic gene subsampling, gene dropout, and, for selected global views, expression masking. The student is therefore trained to recover representations consistent with the teacher despite observing incomplete and perturbed versions of the same cell.

#### Multi-view transcriptomic augmentation

For each cell, all detected genes are first collected and ranked by expression level, and at most the 2,048 highest-expression genes are retained to form the teacher gene pool. The teacher receives a single global view containing all genes in this pool, together with their corresponding expression-bin information. This view provides the most complete transcriptomic observation available to the model during pretraining.

Six student views are independently generated from the same teacher gene pool. Two student global views are constructed by randomly sampling a fraction of genes from the teacher pool:

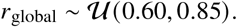

The target number of genes is obtained by rounding *r*_global_*N*_teacher_. When the teacher pool contains more than 1,000 genes, the global-view target size is constrained to be at least 1,000 genes whenever possible; when fewer than 1,000 genes are available, all available genes may be retained. Global views therefore preserve a substantial fraction of the transcriptome while introducing stochastic variation in gene composition.

Four local student views are sampled independently from the same teacher pool. Each local view contains a randomly selected number of genes between 256 and 768:

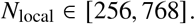

When fewer than 256 genes are available, all available genes are used. Local views therefore provide substantially less transcriptomic information than global views and constitute stronger perturbations of the underlying cell state.

Following subset sampling, independent gene dropout is applied to student views. Each gene in a global student view is dropped with probability 0.10, whereas each gene in a local student view is dropped with probability 0.20. The implementation preserves the configured minimum target size whenever possible, preventing dropout from reducing a view below its minimum retained-gene requirement. No gene dropout is applied to the teacher view.

Student views are sampled independently, and no explicit overlap constraint is imposed between different student views. However, because all student views are sampled from the same teacher gene pool, every retained student gene is guaranteed to occur in the corresponding teacher view. This property enables direct teacher–student matching at the gene level. Gene order is independently shuffled in each view, and expression bins are independently recomputed from the expression values retained in that view.

Together, these augmentations expose scRep to multiple biologically consistent but partially observed versions of the same cell, encouraging the model to capture intrinsic cellular programs rather than memorize individual sparse expression patterns.

#### View-specific expression masking

In addition to gene subsampling and gene dropout, scRep applies expression-specific masking to a subset of student global views. Unlike gene dropout, which removes an entire gene token from the input sequence, expression masking preserves gene identity while hiding only its expression information.

Each cell contributes two student global views. Across all global student views in the distributed mini-batch, 50% are selected for expression masking. For each selected view, a masking ratio is independently sampled as

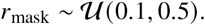

Each retained gene token is then independently selected for masking with probability *r*_mask_. For a masked gene, the gene-identity token remains unchanged, whereas its expression-bin token is replaced by a dedicated [MASK] expression identifier:

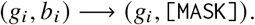

Masked genes remain valid tokens in the Transformer sequence and can therefore interact with all other genes through self-attention.

Expression masking is performed before expression discretization. After the masked-gene set ℳ has been selected, equal-frequency expression binning is computed only from the remaining unmasked genes:

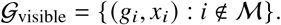

Genes in ℳ do not contribute to the estimation of bin boundaries and are directly assigned the [MASK] expression-bin identifier. This prevents the expression values of masked genes from leaking indirectly through the construction of expression bins. Global student views not selected for masking, all four local student views, and the teacher global view use unmasked expression bins.

This strategy forces the model to infer the state of a masked gene from its identity, the surrounding gene context, and the overall cellular state rather than directly relying on its observed expression value.

#### Cell-level self-distillation objective

For each input view, the final 768-dimensional hidden state of the [CLS] token is used as the cell representation. Rather than directly matching teacher and student [CLS] embeddings in the encoder space, scRep projects these representations into a shared prototype space and performs distribution-level self-distillation.

Let *f*_*θ*_ and *f* _*ξ*_ denote the student and teacher encoders. For a student view *v*_*s*_ and the teacher view *v*_*t*_ from the same cell, the corresponding cell representations are

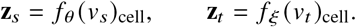

The student and teacher use separate cell-level projection heads with identical architectures. Given a 768-dimensional cell representation, each projection head consists of three bias-free linear layers,

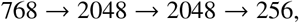

with GELU activations following the first two layers. The resulting 256-dimensional bottleneck representation is normalized by LayerNorm and subsequently ℓ_2_-normalized. A final bias-free prototype layer maps the normalized representation to 4,096 logits.

Let *K* = 4,096 denote the number of learned prototypes. For the teacher global view, the prototype logits are centered and sharpened to construct a stop-gradient target distribution:

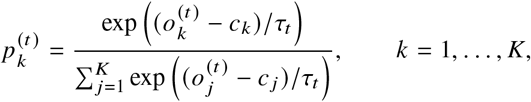

where **o**^(*t*)^ ∈ ℝ^*K*^ denotes the teacher prototype logits, **c** ∈ℝ^*K*^ is the running teacher-output center, and *τ*_*t*_ = 0.04 is the fixed teacher temperature. The teacher distribution is detached from the computational graph.

For each student view *v* ∈ {1, …, 6}, the student prototype logits **o**^(*s,v*)^ ∈ ℝ^*K*^ are converted to temperature-scaled log-probabilities:

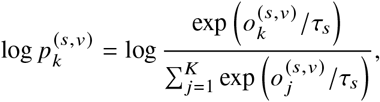

where *τ*_*s*_ = 0.10 is the fixed student temperature. The cell-level self-distillation objective is the mean cross-entropy between the single teacher distribution and the distributions predicted from the six student views:

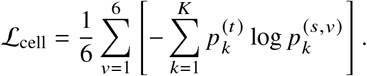

All six student views are therefore weighted equally. By requiring global and local student views to match the same stable teacher target, this objective promotes cell representations that are invariant to gene subsampling, dropout, and expression perturbation.

#### Gene-level self-distillation objective

In addition to cell-level representation learning, scRep performs gene-level self-distillation to directly supervise contextualized gene-token representations. This objective is applied only to student global views selected for expression masking; local student views and unmasked global views do not contribute to the gene-level loss.

For a masked student global view, let ℳ denote the set of genes whose expression-bin information has been masked. For each *g*_*i*_ ∈ℳ, the contextualized student token state 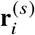 is extracted from the final Transformer layer. The corresponding teacher state 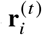 is obtained from the teacher global view of the same cell.

Teacher and student gene representations are aligned by gene identity rather than token position. Specifically, for each masked gene in the student sequence, scRep identifies the token corresponding to the same gene ID in the teacher sequence and forms a matched teacher–student pair. This alignment is required because genes are independently sampled and randomly shuffled across views, such that the same gene generally appears at different sequence positions.

Matched teacher and student gene states are passed through separate gene-level projection heads with independent parameters. These heads have the same architecture as the cell-level projection heads: each 768-dimensional contextualized gene state is mapped to a 256-dimensional normalized bottleneck representation and subsequently projected onto *K* = 4,096 learned prototypes. Thus, *K* denotes the dimensionality of the self-distillation prototype space, rather than the number of genes or gene classes.

For a matched teacher gene state, the resulting prototype logits are centered and sharpened to form a stop-gradient target distribution:

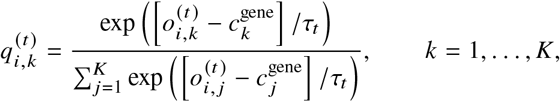

where 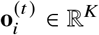 denotes the teacher prototype logits for gene *i*, **c**^gene^ ∈ ℝ^*K*^ is the running gene-level teacher-output center, and *τ*_*t*_ = 0.04 is the fixed teacher temperature. Teacher target distributions are detached from the computational graph.

For the corresponding student gene state, the prototype logits are converted to temperature-scaled log-probabilities:

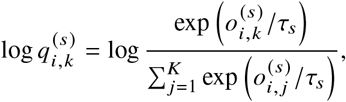

where *τ*_*s*_ = 0.10 is the fixed student temperature. The gene-level objective is computed over all valid matched teacher–student pairs corresponding to masked student genes:

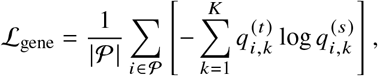

where *P* denotes the set of valid masked-gene teacher–student pairs across eligible student global views.

Only masked genes contribute to this objective; no auxiliary token-level loss is assigned to unmasked student genes. Because the student’s expression information for these genes is hidden while the teacher observes the complete expression state, the student must infer the teacher’s contextualized representation from gene identity, neighboring genes, and cell-level context. This encourages scRep to capture context-dependent gene activity and co-expression relationships rather than only static gene identity.

#### Overall training objective and teacher update

The overall pretraining objective combines cell-level and gene-level self-distillation:

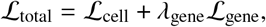

With

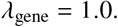

Thus, the two objectives are given equal weight. Only the student Transformer backbone, student cell-level projection head, and student gene-level projection head receive gradients. The teacher backbone and both teacher projection heads are initialized from the student and subsequently updated by EMA after every optimizer step:

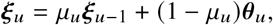

where *u* denotes the optimization step. The teacher momentum follows a cosine schedule from *μ*_0_ = 0.996 at the beginning of training to *μ*_end_ = 1.0 at the final training step:

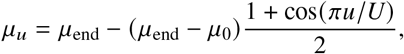

where *U* is the total number of optimization steps. Increasing the teacher momentum toward one makes the teacher progressively more stable during later stages of training.

Teacher output centers are updated after each forward pass using the distributed mean of teacher logits. For the cell-level projection head,

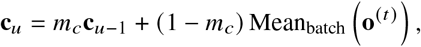

where *m*_*c*_ = 0.9. The gene-level center is updated analogously from teacher logits corresponding to matched masked-gene pairs, with *m*_gene_ = 0.9. Batch statistics are aggregated across distributed training workers before updating the centers.

The teacher temperature is fixed at *τ*_*t*_ = 0.04, whereas the student temperature is fixed at *τ*_*s*_ = 0.10, with no temperature warm-up. The lower teacher temperature produces sharper target distributions, while teacher centering helps prevent collapse toward a small subset of prototype dimensions. Both cell-level and gene-level losses are averaged over their valid teacher–student pairs, preventing views or cells with more retained genes from disproportionately dominating the training objective.

After pretraining, the momentum teacher encoder is used as the default feature extractor for downstream analyses. The final [CLS] hidden state is used as the cell-level representation, whereas contextualized gene-token hidden states provide gene-level representations.

### 4.4 Datasets and data processing

#### CELLxGENE scRNA-seq and snRNA-seq collection

We collected human single-cell and single-nucleus transcriptomic data from the CELLxGENE portal (https://cellxgene.cziscience.com/) using the CELLxGENE Census API (https://chanzuckerberg.github.io/cellxgene-census/python-api.html). We used the Census release dated November 8, 2025, and included both scRNA-seq and snRNA-seq datasets. To remove low-complexity cells and nuclei, we retained profiles with more than 1,000 detected unique genes and more than 3,000 total counts.

From the quality-controlled collection, we first identified datasets eligible for independent downstream evaluation. We then selected six benchmark datasets spanning different tissues, disease contexts, sequencing modalities, and dataset sizes, with the specific datasets chosen from eligible candidates without reference to downstream model performance.

All six benchmark datasets were excluded from the scRep pretraining corpus to prevent overlap between pretraining and downstream evaluation.

The resulting CELLxGENE pretraining pool contained 59,835,009 eligible profiles. The large-scale scRep model was trained on 30.72 million distinct profiles sampled from this pool using a global mini-batch size of 512 profiles. The checkpoint analyzed in this study was obtained after 60,000 optimization steps, corresponding to the processing of these 30.72 million profiles. We therefore refer to this model as scRep-30M.

#### Tabula Sapiens human cell atlas

We obtained the Tabula Sapiens human cell atlas from the Tabula Sapiens data portal [7]. Tabula Sapiens is a cross-tissue atlas of healthy adult human cells spanning multiple organs and major immune, epithelial, endothelial, stromal, muscular, and neural compartments, generated using both 10x Genomics and Smart-seq2 technologies.

We used the Tabula Sapiens expression data as a human reference corpus for self-supervised pretraining. The downloaded data were converted to AnnData format, harmonized to the shared 19,238-gene vocabulary, and partitioned into shards for streaming training. No cell-type, tissue, donor, or other metadata labels were used as model supervision; each cell contributed only its gene-expression profile to the self-distillation objective.

#### Human posterior ocular segment transcriptome

We obtained the human posterior ocular-segment dataset from the CZ CELLxGENE Discover collection *Transcriptomic Analysis of the Ocular Posterior Segment Completes a Cell Atlas of the Human Eye* (dataset ID: `9f1a899f-05a0-44c5-8aa9-39df61ae4324`; https://cellxgene.cziscience.com/collections/7855daa4-3f34-4aae-b65a-87720c02e7cd). The dataset contains 139,978 cells from 19 donors profiled using 10x Genomics 3^′^ v3 and spans multiple posterior-eye tissues, including retina, retinal pigment epithelium, macula, optic disc, optic nerve, and sclera.

#### Human pancreatic islet atlas

We downloaded the pancreatic-islet dataset from the CZ CELLxGENE Discover collection *A pancreatic islet scRNA-seq atlas from 48 non-diabetic, prediabetic and type 2 diabetic individuals of matched demographic and ethnic backgrounds* (dataset ID: e51bae9a-c747-4b64-904a-4da7cda218ab; https://cellxgene.cziscience.com/collections/58e85c2f-d52e-4c19-8393-b854b84d516e). The dataset contains 245,878 cells from 48 donors, including non-diabetic, prediabetic, and type 2 diabetic individuals, profiled using 10x Genomics 3^′^ v2 and v3 assays and spanning major endocrine, exocrine, stromal, vascular, and immune populations.

#### Human brain vascular single-nucleus multi-omics dataset

We obtained the single-nucleus RNA-sequencing component of the human brain vascular multi-omics dataset from CZ CELLxGENE Discover (dataset ID: 203025fe-fa99-4d57-81da-458ed8f0c334; https://cellxgene.cziscience.com/collections/433700dc-e8a5-48b0-b5cd-beb22f3f88fe). The dataset, entitled *Brain vascular single-cell multi-omics elucidates disease risk associations*, contains 65,479 nuclei from dorsolateral prefrontal cortex samples from 30 donors, including individuals with Alzheimer disease, cognitive disorder, and no reported disease. The RNA modality was generated using the 10x Genomics Multiome platform.

#### Glioblastoma single-cell RNA-sequencing dataset

We downloaded the glioblastoma dataset from the CZ CELLxGENE Discover collection *Single-cell RNA sequencing of glioblastoma cases with tissue sampled from tumor core to macroscopically normal cortex* (dataset ID: 145dcf6a-2461-4fa3-a0af-7fc56db0bd33; https://cellxgene.cziscience.com/collections/113a558a-e96e-4643-81db-140e95c58578). We used the SL057 sample, containing 123,236 cells from the right temporal lobe of a 63-year-old male donor with glioblastoma, profiled using 10x Genomics 3^′^ v3 chemistry. The dataset spans malignant, immune, and brain-resident cell populations.

#### Human kidney with ureteral obstruction

We obtained the kidney dataset from the CZ CELLxGENE Discover collection *snRNA-seq of human kidney with ureteral obstruction* (dataset ID: 867757c1-3b1a-49d9-a0cd-17767eb160cc; https://cellxgene.cziscience.com/collections/307ce143-7cc8-4813-99b1-3797834149c9). The dataset contains 46,957 nuclei from 12 donors with either normal kidney tissue or obstructive nephropathy and was generated using the 10x Genomics Multiome platform. It includes major tubular, endothelial, stromal, and immune cell populations.

#### Human myocarditis cardiac-tissue dataset

We downloaded the cardiac dataset from the CZ CELLxGENE Discover collection *The cellular and molecular cardiac tissue responses in human inflammatory cardiomyopathies after SARS-CoV-2 infection and COVID-19 vaccination* (dataset ID: fe7aae33-6f7c-41a5-8d29-9996a9ddf1ab; https://cellxgene.cziscience.com/collections/328d71f0-0ed7-4518-966f-be6bd0797324). Our standardized evaluation subset contains 62,705 cells from the left ventricle of 24 donors, including post-COVID-19, post-vaccination, multisystem inflammatory syndrome, and non-COVID-19 control samples. Cells were profiled using 10x Genomics 3^′^ v3 chemistry.

#### GSE109555: human peri-implantation embryonic development

We downloaded GSE109555 from the NCBI Gene Expression Omnibus (GEO; https://www.ncbi.nlm.nih.gov/geo/query/acc.cgi?acc=GSE109555). This study, entitled *Reconstituting the transcriptome and DNA methylome landscapes of human implantation*, profiled more than 8,000 cells from 65 human peri-implantation embryos using single-cell multi-omics assays. We used the released transcriptomic matrix, `GSE109555_All_Embryo_TPM.txt.gz`, for trajectory analysis.

The dataset contains embryonic cells collected across developmental days 6–14. Gene symbols were mapped to the scRep gene vocabulary, and cells were encoded using the teacher backbone. We constructed a *k*-nearest-neighbor graph from scRep embeddings, performed Leiden clustering, computed PAGA connectivity, and estimated diffusion pseudotime. The trajectory root was selected from cells from the earliest developmental day, and agreement between inferred pseudotime and collection day was quantified by Spearman correlation. Collection-day labels were used only as an external temporal reference and were not used to construct the embedding graph or train the model.

#### GSE117498: human hematopoietic progenitor differentiation

We downloaded GSE117498 from GEO (https://www.ncbi.nlm.nih.gov/geo/query/acc.cgi?acc=GSE117498). This study, entitled *A comprehensive single cell transcriptional landscape of human hematopoietic progenitors*, contains single-cell RNA-sequencing profiles of lineage-negative human bone-marrow hematopoietic stem and progenitor cells, including HSCs, MPPs, MLPs, pre-B/NK progenitors, CMPs, GMPs, and MEPs.

We concatenated the raw-count matrices from the FACS-sorted populations, mapped gene symbols to the scRep gene vocabulary, and embedded individual cells using the teacher backbone. A *k*-nearest-neighbor graph, Leiden clustering, PAGA lineage graph, and diffusion pseudotime were inferred solely from scRep representations. The trajectory root was selected using an HSC-marker score based on *HLF, AVP, MEIS1, GATA2, PROCR, MPL*, and *MLLT3*. FACS population labels were withheld during embedding and trajectory inference and used only for external validation of the inferred pseudotime ordering and lineage structure.

### 4.5 Pre-training corpus construction

We constructed five human single-cell pre-training corpora to examine the effects of training-set scale and cell-type diversity. The corpora were based on Tabula Sapiens, CELLxGENE, or combinations of the two.

The large-scale CELLxGENE corpus used for scRep-30M was drawn from the quality-controlled CELLxGENE pretraining pool described above, for which cells were required to have more than 1,000 detected genes and more than 3,000 total counts. For construction of the smaller Tabula Sapiens–CELLxGENE scaling corpora, we applied an additional, more stringent cell-complexity criterion: candidate CELLxGENE cells were required to have `n_genes_by_counts ¿` 2,000 and a valid `cell_type_ontology_term_id`. This additional criterion was used only for the controlled smaller-scale corpus construction and was not applied to the scRep-30M corpus.

The six downstream benchmark datasets were excluded from all CELLxGENE-derived pretraining corpora to prevent train–test overlap. Corpus construction and cell selection were performed independently of downstream evaluation results.

#### Tabula Sapiens 1.13M

The reference corpus contained all 1,136,218 Tabula Sapiens cells and 180 observed cell types. This corpus was used as the baseline for the nested scaling analysis.

#### Tabula Sapiens + CELLxGENE 2M

We retained all 1,136,218 Tabula Sapiens cells and added 863,782 quality-filtered CELLxGENE cells, resulting in a corpus of exactly 2,000,000 cells and 853 observed cell types. CELLxGENE cells were selected using a coverage-first strategy based on `cell_type_ontology_term_id`. Specifically, ontology terms absent from Tabula Sapiens were prioritized, followed by rare Tabula Sapiens cell types. For each ontology term, the target abundance depended on the number of eligible CELLxGENE candidates: terms with fewer than 100, 100–499, 500–1,999, 2,000–9,999, and at least 10,000 available cells were capped at 100, 300, 1,000, 2,000, and 5,000 cells, respectively. Existing Tabula Sapiens types were supplemented only up to these targets, whereas absent types could receive the full target. Allocation first aimed to provide each eligible type with up to 300 cells and then distributed the remaining budget toward the type-specific cap. The supplement included cells from 553 CELLxGENE source datasets, 297 tissue ontology terms, and 720 sampled ontology terms. Overall, 673 cell types were newly introduced relative to Tabula Sapiens.

#### Tabula Sapiens + CELLxGENE 2.8M

We applied the same coverage-first selection procedure with a larger sampling budget. This corpus contained all 1,136,218 Tabula Sapiens cells plus 1,667,317 CELLxGENE cells, yielding 2,803,535 cells in total and 853 observed cell types. Although this corpus is referred to as “3M” during dataset generation, its final size was 2.80M cells because the available eligible cells and per-cell-type caps did not support a full 3M-cell corpus.

The CELLxGENE supplement was drawn from 579 source datasets, 324 tissue ontology terms, and 773 sampled ontology terms. As in the 2M corpus, it introduced 673 cell types not present in Tabula Sapiens.

#### Balanced 2M subset

To distinguish the influence of cell-type composition from that of total cell number, we constructed a 2,000,000-cell subset from the 2.8M Tabula Sapiens + CELLxGENE corpus. Cells were grouped by `cell_type_ontology_term_id`. Cell types with no more than 5,000 cells were retained completely. For larger cell types, an initial cap of 5,000 cells per type was imposed and then reduced approximately equally across the large cell types until the total reached 2,000,000 cells. This affected 42 large cell types, which received final quotas of 4,692 or 4,693 cells. Within each cell type, cells with `n_genes_by_counts` ≥ 2,000 were retained preferentially, with lower-depth cells included only if needed to meet the quota. The final corpus contained 2,000,000 cells and retained all 853 observed cell types; it comprised 332,683 Tabula Sapiens cells and 1,667,317 CELLxGENE-derived cells.

#### CELLxGENE 30.72M

The large-scale CELLxGENE training corpus was constructed from the quality-controlled human CELLxGENE pretraining pool after exclusion of the six held-out evaluation datasets. This pool contained approximately 59.8 million eligible transcriptomic profiles after applying the initial collection-level quality-control filters of more than 1,000 detected genes and more than 3,000 total counts. From this pool, 30.72 million distinct profiles were used to train scRep-30M. Training used a global mini-batch size of 512 profiles, and the checkpoint analyzed in this study was obtained after 60,000 optimization steps, corresponding to processing the full 30.72-million-profile training set. The additional `n_genes_by_counts` > 2,000 criterion used for construction of the smaller Tabula Sapiens–CELLxGENE scaling corpora was not applied to this large-scale training corpus.

The resulting 30.72-million-profile corpus comprised data from 833 source datasets and 856 observed cell types. Unlike the hybrid Tabula Sapiens–CELLxGENE corpora, this corpus was not constructed using the coverage-first rebalancing strategy and did not include Tabula Sapiens cells. It was therefore treated as an external large-scale training composition rather than as part of the nested Tabula Sapiens expansion series.

### 4.6 Pre-training corpus diversity analysis

We quantified the diversity of each pre-training corpus from its observed cell-type abundance distribution. For a corpus containing *K* cell types and *n*_*i*_ cells assigned to cell type *i*, we defined the abundance-aware diversity index as

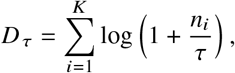

where *τ* is an abundance-scale parameter. The logarithmic transformation imposes diminishing marginal contributions as additional cells are added to an already represented cell type, thereby reducing the dominance of highly abundant populations while retaining contributions from rare and moderately represented cell types. The parameter *τ* controls the abundance scale at which this saturation occurs: smaller values cause abundant cell types to saturate more rapidly, whereas larger values retain greater sensitivity to within-type cell abundance. Thus, *D* _*τ*_ reflects both cell-type richness and the distribution of cell abundance across represented cell types.

The abundance-scale parameter *τ* was selected using only the nested Tabula Sapiens expansion series: Tabula Sapiens 1.13M, TS+CXG 2M, and TS+CXG 2.8M. Their corresponding mean clustering performance values were 0.760987, 0.824935, and 0.831610, respectively, with performance summarized using the evaluation protocol described below.

We optimized *τ* over the interval [10^−18^, 10^6^] on a logarithmic scale. For each candidate value of *τ*, we calculated *D* _*τ*_ for the three corpora and fitted an ordinary least-squares linear relationship,

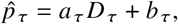

where 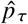 denotes predicted mean clustering performance. We selected the value of *τ* that minimized the root mean squared error between predicted and observed performance across the three nested corpora. This procedure yielded

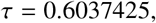

with the fitted relationship

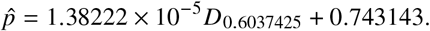

All diversity values reported in Fig. 3 were calculated directly from *D*_0.6037425_; the fitted slope and intercept were used only to summarize the diversity–performance relationship and were not used to transform the reported diversity scores. The trend line shown in Fig. 3 was fitted using the same three nested scaling corpora.

The Balanced 2M and CELLxGENE 30.72M compositions were excluded from both *τ* optimization and trend-line fitting and were instead treated as validation compositions. Balanced 2M was derived from the TS+CXG 2.8M corpus by reducing the abundance of highly represented cell types while retaining all observed cell types, thereby providing a matched-scale composition comparison. The CELLxGENE 30.72M composition represented a substantially larger, independently composed training corpus outside the nested Tabula Sapiens–CELLxGENE scaling series.

Using *τ* = 0.6037425, the abundance-aware diversity scores were 1,290.94 for Tabula Sapiens 1.13M, 5,917.39 for TS+CXG 2M, 6,400.35 for TS+CXG 2.8M, 6,351.41 for Balanced 2M, and 7,103.87 for CELLxGENE 30.72M. The corresponding numbers of observed cell types were 180, 853, 853, 853, and 856, respectively.

Because *τ* was selected using only three nested corpora, the fitted relationship should be interpreted as a descriptive summary of the observed diversity–performance trend rather than as evidence that *τ* represents a uniquely identifiable biological diversity scale.

### 4.7 Baseline foundation models

We compared scRep with six publicly available single-cell foundation models: Geneformer-v2, scGPT, scFoundation, scConcept, STACK, and STATE. For every baseline, we used the released pretrained checkpoint in inference mode and did not fine-tune the model on any evaluation dataset. Gene identifiers were mapped to the vocabulary required by each model; genes without a valid mapping were omitted. We extracted one embedding per cell using the model-specific embedding procedure described below. The resulting cell embeddings were used as the input to the common downstream evaluation pipeline.

#### Geneformer-v2

We used the released Geneformer-v2 104M-parameter checkpoint (`Geneformer-V2-104M`) [39, 5]. Geneformer represents a cell as an ordered sequence of genes ranked by within-cell expression, thereby emphasizing relative transcriptional priority rather than absolute expression magnitude. Input sequences were truncated to the Geneformer-v2 maximum length of 4,096 tokens. We used the pretrained model without task-specific fine-tuning and extracted the final-layer cell representation using its standard cell-embedding interface.

#### scGPT

We evaluated the released pretrained scGPT checkpoint [8]. scGPT is a transformer model that jointly encodes gene tokens and discretized gene-expression values. Following the official cell-embedding workflow, we mapped input genes to the scGPT vocabulary, used a maximum sequence length of 1,200 tokens including the `<cls>` token, and extracted the `<cls>` representation as the cell embedding. Embeddings were ℓ_2_-normalized before downstream analysis.

#### scFoundation

We used the released scFoundation pretrained model and its official embedding workflow [15]. scFoundation is a gene-expression foundation model trained with masked expression reconstruction objectives. We mapped genes to the model’s 19,264-gene reference index and encoded the positive-expression genes in each cell. Cell embeddings were obtained by the official all-token pooling procedure (rather than max pooling) and were ℓ_2_-normalized before evaluation.

#### scConcept

We evaluated the released scConcept checkpoint [2]. scConcept uses a contrastive representation-learning framework that combines gene identity and expression information to generate cell embeddings. Genes were mapped to the human scConcept vocabulary; for each cell, the top nonzero-expression genes were supplied to the model, with at most 20,000 input tokens. We enabled the model’s learned gene embeddings, extracted the standard cell-level embedding, and applied ℓ_2_ normalization.

#### STACK

We used the released STACK-Large checkpoint (`bc_large.ckpt`) [12]. STACK is a 217M-parameter encoder–decoder model that uses tabular attention, alternating gene-wise and cell-wise attention over cell-by-gene matrix chunks. The released STACK-Large model was pretrained for 10 epochs on approximately 150 million uniformly processed human single cells. We mapped input genes to the released STACK base-count gene list and used the pretrained model to extract cell embeddings without fine-tuning.

#### STATE

We used the State Embedding (SE)-600M model from the STATE framework [1]. STATE combines expression measurements with pretrained gene/protein embeddings to generate cell-state representations. Specifically, we used the released SE-600M checkpoint after 16 training epochs (`se600m epoch16.ckpt`), together with its matched protein-embedding file and configuration. Inputs were processed through the official STATE embedding transformation, and the resulting cell embeddings were used without task-specific fine-tuning.

### 4.8 Evaluation protocols

Unless otherwise stated, all analyses used frozen pretrained encoders and no task-specific fine-tuning on the evaluation data. We used the same held-out human test datasets, cell identifiers, and ground-truth metadata for every method within a comparison. All stochastic sampling and graph-based analyses used a fixed random seed.

#### 4.8.1 Cell clustering and cell-type annotation

##### Held-out cell-type datasets and embeddings

We evaluated representations on six held-out human single-cell datasets from diverse tissues and disease contexts: glioblastoma, brain, kidney, eye, pancreas, and heart. The datasets were excluded during construction of the CELLxGENE supplements used for scRep pre-training. For each encoder, we computed one cell embedding per cell and aligned representations across methods by the unique cell identifier. Cells without a valid `cell_type` label were excluded. Thus, each method was evaluated on exactly the same labeled cells within a dataset.

##### Unsupervised cell clustering

We evaluated clustering using the complete cell embedding without a preliminary PCA reduction or feature standardization. For each dataset and method, we constructed a 30-nearest-neighbor graph using Euclidean distance in embedding space (Scanpy neighbors; random seed 42), followed by Leiden community detection implemented through EpiScanpy. The Leiden resolution was automatically adjusted with getNClusters until the number of inferred clusters matched the number of annotated cell types in that dataset. This calibration controls for trivial differences caused solely by a method producing more or fewer clusters. Predicted cluster assignments were compared with reference cell-type labels using adjusted Rand index (ARI) and normalized mutual information (NMI), with arithmetic NMI normalization. The primary clustering score was the mean of ARI and NMI. Scores were computed separately for each of the six datasets and then averaged with equal dataset weight; no pooled-cell metric was used. UMAP visualizations were calculated from the same 30-nearest-neighbor graph and were used only for visualization, not for metric calculation.

##### Reference-based cell-type annotation

We assessed label transfer with a group-disjoint k-nearest-neighbor (kNN) protocol. For each dataset, an informative donor, patient, sample, batch, or library identifier was selected as the grouping variable, in that order of availability. Groups were randomly permuted and divided by five-fold cross-validation; each fold used cells from 20% of groups as a reference pool and cells from the remaining 80% of groups as the query set. This prevents the same donor/sample group from contributing both reference and query cells.

Within each reference pool, cells were sampled independently per cell type at reference fractions of 0.5%, 1%, 2%, 5%, 10%, and 20%, with a minimum of one and a maximum of 100 reference cells per cell type. We additionally evaluated the full available reference pool. A cell type was evaluated in a fold only if it occurred in both reference and query cells and had at least one query cell; folds containing fewer than 20 query cells were excluded. Query labels were predicted from the 20 nearest reference embeddings using cosine similarity. Neighbor votes were weighted by softmax (*s*/ 0.07), where *s* is cosine similarity. We report macro-F1 as the annotation metric because it gives equal weight to common and rare cell types. Macro-F1 was first calculated within each group-disjoint fold and then summarized across folds and datasets.

#### 4.8.2 Pre-training scaling evaluation

We compared five pre-training settings: Tabula Sapiens 1.13M (1,136,218 cells; 180 observed cell types), TS+CXG 2M (2,000,000 cells; 853 cell types), TS+CXG 2.8M (2,803,535 cells; 853 cell types), Balanced 2M (2,000,000 cells; 853 cell types), and the large-scale CELLxGENE training setting used for scRep-30M.

The first three corpora formed a nested Tabula Sapiens–CELLxGENE scaling series. Balanced 2M was a cell-type-balanced downsample of TS+CXG 2.8M and served as a matched-scale composition comparison. In contrast, scRep-30M was trained on 30.72 million distinct profiles sampled from the 59.8-million-profile CELLxGENE pre-training pool and was treated as an external large-scale setting rather than as part of the nested scaling series.

Pre-training corpus diversity was quantified using the abundance-aware diversity index described in the Pre-training corpus diversity analysis section.

For each pre-training run, downstream clustering performance was evaluated on six held-out benchmark datasets— Glioblastoma, Brain, Kidney, Eye, Pancreas, and Heart—comprising 684,233 cells in total. Performance was summarized at three pre-specified late-training checkpoints at 55k, 60k, and 65k training steps. These checkpoints were separated by 5,000 training steps to represent distinct model states rather than closely spaced snapshots.

At each checkpoint, calibrated ARI and calibrated NMI were computed for each of the six benchmark datasets, yielding 12 dataset–metric measurements. Checkpoint-level performance was defined as the arithmetic mean of these 12 values, and run-level performance was obtained by averaging the three checkpoint-level scores. This fixed checkpoint window was used instead of selecting the single best-performing checkpoint in order to reduce sensitivity to individual model snapshots.

For scRep-30M, the 30.72-million-profile composition encountered through the 60k checkpoint was used as the representative large-scale composition for diversity calculation, while downstream performance was summarized across the same pre-specified 55k–65k checkpoint window used for the smaller-scale runs.

#### 4.8.3 Gene-level analyses

##### Marker-gene sensitivity in the Eye dataset (Fig. 4a–c)

We evaluated whether scRep representations exhibit elevated sensitivity to established cell-type marker genes in the Eye test dataset using the frozen scRep TS+3M checkpoint at 65,000 training steps. Each cell was represented by its top 2,048 expressed genes, with expression values discretized into 50 bins. We analyzed cell types containing at least 20 cells and retained genes detected in at least 2% of cells within the corresponding cell type.

For each eligible cell–gene instance, we retained the gene identity token and all other model inputs but replaced the expression-bin channel of the selected gene with the model padding bin. Let **z** and **z**_−*g*_ denote the original and perturbed ℓ_2_-normalized CLS embeddings, respectively. We defined the per-instance perturbation effect as

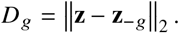

For each gene, perturbation effects were averaged over cells in which that gene was observed, yielding the conditional mean L2 mask effect. Genes were ranked within each cell type according to this quantity and converted to within-cell-type percentiles. Figure 4a compares the median conditional-L2 percentile of curated marker genes with the median percentile of all other eligible genes.

To account for the frequency with which a gene contributes across the cell population, we additionally defined a population-level mask effect as the conditional mean L2 mask effect multiplied by the gene’s detection frequency within that cell type. Figure 4b reports the corresponding within-cell-type percentile ranks. Thus, the conditional score measures representation sensitivity given that a gene is observed, whereas the population-level score additionally accounts for the proportion of cells in which the gene can contribute.

For Fig. 4c, each marker gene was matched to 50 non-marker genes from the same cell type using nearest-neighbor matching based on standardized detection frequency and standardized log (1 + mean expression). For each cell type, we report the fraction of marker genes whose conditional-L2 percentile exceeded the median percentile of their matched controls. The Eye analysis comprised 16 cell types and 147 cell-type–marker-gene pairs represented in both the dataset and the model vocabulary.

These perturbation scores quantify the sensitivity of the learned cell representation to the observed expression state of individual genes and should not be interpreted as causal feature attributions.

##### Contextual gene embeddings and TF-associated regulons in the Glioblastoma dataset (Fig. 4d–f)

Panels d–f were generated from the Glioblastoma test dataset using the same frozen scRep checkpoint at 65,000 training steps. Contextualized token representations for each gene were averaged across all cells in which that gene was observed to obtain a dataset-level gene embedding, after which the resulting embeddings were ℓ_2_-normalized.

We constructed a dataset-specific directed similarity network using curated human transcription factors as source nodes. For each TF, candidate target genes were ranked by cosine similarity to the TF embedding, and up to 20 targets with cosine similarity greater than 0.2 were retained. Edge weights were defined by the corresponding cosine similarities. Figure 4d shows the ten highest-weight candidate targets for SOX5 and SPI1.

This procedure defines an embedding-similarity network and should be interpreted as identifying putative TF-associated gene programs rather than validated direct regulatory interactions or supervised GRN predictions.

For Fig. 4e–f, each TF-associated target set was treated as an embedding-derived regulon. The regulon score for each cell was calculated as the edge-weighted mean log (1 + *x*) expression of its retained target genes, with TF–target cosine similarities used as edge weights.

For the paired heatmaps in Fig. 4e, regulon scores and TF expression were averaged within cell types. Cell types containing fewer than 50 cells were excluded, as were unknown and neoplastic-cell labels. Values for each TF were then standardized across cell types to facilitate comparison of lineage-specific activity patterns.

Figure 4f projects cell-type annotations, SOX5 and SPI1 regulon scores, and the corresponding TF expression values onto a shared UMAP representation of the Glioblastoma cells. These analyses compare embedding-derived regulon activity with TF expression within the same dataset and do not treat the inferred TF–target similarities as experimentally validated regulatory relationships.

#### 4.8.4 Trajectory analyses

##### Embryonic-development trajectory

We evaluated developmental trajectory preservation using the GSE109555 human peri-implantation embryo expression dataset. Gene symbols were mapped to Ensembl identifiers, duplicated mapped genes were removed, and cells were embedded using the frozen scRep model. For each cell, the CLS embedding was obtained from the top 2,048 expressed genes using 50 expression bins.

A 30-nearest-neighbor graph was constructed in the scRep embedding space. UMAP coordinates were generated for visualization, Leiden clustering was performed at a resolution of 0.8, PAGA connectivity was calculated, and diffusion components were computed. Diffusion pseudotime (DPT) was then inferred from a single early-state root [14].

To orient the trajectory, the root was selected as the medoid in scRep embedding space among cells sampled at the earliest developmental day (D6). Developmental-day labels were not used for embedding generation, neighborhood-graph construction, Leiden clustering, PAGA connectivity, diffusion-map calculation, or the geometry of the DPT trajectory; they were used only to restrict root selection to the earliest sampled population and subsequently to evaluate temporal agreement.

Temporal concordance was quantified using the Spearman correlation between developmental day and inferred pseudotime across cells.

##### CD34+ bone-marrow differentiation

We evaluated haematopoietic trajectory preservation using the GSE117498 human CD34+ bone-marrow dataset, which contains experimentally defined FACS populations spanning haematopoietic stem and progenitor states. Gene symbols were mapped to Ensembl identifiers, duplicated identifiers were removed, and cells were embedded using frozen scRep with the top 2,048 expressed genes and 50 expression bins.

A 30-nearest-neighbor graph was constructed in embedding space, followed by UMAP visualization, Leiden clustering at a resolution of 0.6, PAGA connectivity analysis, diffusion-map calculation, and DPT inference. FACS population labels were withheld from all stages of trajectory construction.

The root was selected independently of FACS annotations using an HSC marker-expression score, with the highest-scoring early-state cell used to orient the trajectory. FACS population labels were examined only after pseudotime inference to assess the correspondence between inferred developmental progression and experimentally defined cell states.

As a summary measure of early-to-late ordering, we computed the area under the receiver operating characteristic curve (AUROC) using inferred pseudotime to distinguish HSCs from lineage-restricted progenitors comprising GMP, MEP, and PreB/NK cells. This AUROC was calculated directly from the inferred pseudotime ranking and does not represent the performance of a supervised classifier trained on FACS labels.

### 4.9 Implementation details

scRep was implemented as a self-distillation transformer for single-cell gene expression. The encoder comprised 12 transformer blocks with 12 attention heads, an embedding dimension of 768, and a feed-forward expansion dimension of 3,072. A CLS token was prepended to each cell sequence and its final representation was used as the cell embedding. Student and teacher networks had the same architecture; the teacher weights were updated as an exponential moving average of the student weights. The projection head used for self-distillation had an output dimension of 4,096, a hidden dimension of 2,048, and a bottleneck dimension of 256.

Only non-zero-expression genes were supplied as input. Their expression values were encoded by value-aware, approximately equal-frequency bins. The teacher view contained up to the 2,048 most highly expressed genes from each cell. To create student views, we generated two global views and four local views through random gene subsampling and gene dropout; local views retained at least 256 genes when that many genes were available. For additional occlusion-based augmentation, half of the global student views in each batch were selected for expression masking, and 10–50% of their gene-expression tokens were masked. Masked positions were excluded before bin construction, preventing their expression values from contributing to the bin labels visible to the student.

The model was optimized with a cell-level self-distillation objective between student and teacher CLS representations together with an equally weighted masked token-level distillation objective. No auxiliary expression-reconstruction objective was used. Thus, training encouraged agreement between differently masked or subsampled views of the same cell at both cell and gene-token levels, without reconstructing the original expression profile. We used AdamW with an initial learning rate of 5 × 10^−5^ and weight decay of 0.01. Each optimization step used a global mini-batch of 512 cells aggregated across all GPUs. The learning rate was linearly warmed up for the first 2,000 optimization steps and then followed a cosine decay schedule to 10% of its initial value over the scheduled training run. Training duration and checkpoint selection followed the run-specific training configuration.

We implemented model training and inference in PyTorch. AnnData objects were handled with anndata and Scanpy. Scanpy was used to construct neighborhood graphs and to compute UMAP, diffusion maps, PAGA, and diffusion pseudotime; the EpiScanpy implementation of Leiden clustering was used for the calibrated clustering evaluation. ARI, NMI, classification metrics, and regression metrics were calculated with scikit-learn.

## Code availability

The source code used to train and evaluate scRep, together with the scripts required to reproduce the main experiments and analyses in this study, will be made publicly available upon publication of the article. The pretrained scRep model checkpoints and associated configuration files will also be released at that time.

## Data availability

All datasets used in this study are derived from publicly available single-cell transcriptomic resources. The corresponding data sources, accession identifiers, and dataset information are described in the Methods section. Processed data and metadata required to reproduce the analyses presented in this study will be made publicly available upon publication of the article, subject to the access conditions and licensing requirements of the original data sources.

---

**Table S1:** Comparison of the pretrained models used in this study.

| Model | Parameters | Pre-training cells |
| --- | --- | --- |
| **scRep-3M** | **99.8M** | **2.804M** |
| **scRep-30M** | **99.8M** | **30.72M** |
| Geneformer-v2 | 104M | ~104M |
| scGPT (whole-human) | 51M | 33M |
| scFoundation | 100M | >50M |
| scConcept | 30M | 40M |
| STACK-Large | 217M | ~150M |
| STATE (SE-600M) | 715M | 167M |

**Notes.** The scRep-3M and scRep-30M entries refer to the two models trained in this study. Their parameter counts are identical because they use the same architecture. For scRep 3M, the planned 3M-cell corpus yielded 2,803,535 eligible cells after quality filtering and coverage-balanced selection. The values for Geneformer-v2, scGPT, scFoundation, scConcept, STACK, and STATE correspond to the released checkpoints used in our experiments. Pre-training cell counts reported as approximate values follow the corresponding model release documentation.

**Table S2:** Per-dataset biological representation quality. Macro-F1 was evaluated using KNN cell-type annotation with the full reference set and is reported as mean ± s.d. across five group-disjoint cross-validation folds. NMI and ARI are obtained from Leiden-calibrated clustering.

| Dataset | Model | NMI ↑ | ARI ↑ | Macro-F1 ↑ |
| --- | --- | --- | --- | --- |
| Glioblastoma | scRep-30M | 0.961 | 0.973 | $0.860 \pm 0.091$ |
| | scRep-3M | 0.944 | 0.957 | $0.840 \pm 0.100$ |
| | STATE (SE-600M) | 0.939 | 0.949 | $0.849 \pm 0.090$ |
| | Stack | 0.887 | 0.800 | $0.841 \pm 0.100$ |
| | scConcept | 0.829 | 0.617 | $0.808 \pm 0.106$ |
| | scFoundation | 0.880 | 0.867 | $0.829 \pm 0.088$ |
| | scGPT | 0.733 | 0.465 | $0.767 \pm 0.096$ |
| | Geneformer-v2 | 0.623 | 0.317 | $0.582 \pm 0.121$ |
| Brain | scRep-30M | 0.846 | 0.763 | $0.925 \pm 0.004$ |
| | scRep-3M | 0.841 | 0.760 | $0.926 \pm 0.004$ |
| | STATE (SE-600M) | 0.757 | 0.577 | $0.879 \pm 0.019$ |
| | Stack | 0.849 | 0.731 | $0.935 \pm 0.003$ |
| | scConcept | 0.801 | 0.699 | $0.935 \pm 0.007$ |
| | scFoundation | 0.710 | 0.508 | $0.932 \pm 0.004$ |
| | scGPT | 0.699 | 0.557 | $0.797 \pm 0.028$ |
| | Geneformer-v2 | 0.668 | 0.476 | $0.865 \pm 0.030$ |
| Kidney | scRep-30M | 0.910 | 0.894 | $0.867 \pm 0.019$ |
| | scRep-3M | 0.890 | 0.879 | $0.870 \pm 0.028$ |
| | STATE (SE-600M) | 0.919 | 0.924 | $0.756 \pm 0.028$ |
| | Stack | 0.863 | 0.790 | $0.826 \pm 0.032$ |
| | scConcept | 0.787 | 0.588 | $0.831 \pm 0.028$ |
| | scFoundation | 0.751 | 0.552 | $0.761 \pm 0.042$ |
| | scGPT | 0.586 | 0.431 | $0.519 \pm 0.046$ |
| | Geneformer-v2 | 0.669 | 0.450 | $0.715 \pm 0.062$ |
| Eye | scRep-30M | 0.914 | 0.874 | $0.914 \pm 0.036$ |
| | scRep-3M | 0.918 | 0.879 | $0.925 \pm 0.032$ |
| | STATE (SE-600M) | 0.905 | 0.864 | $0.887 \pm 0.059$ |
| | Stack | 0.902 | 0.835 | $0.908 \pm 0.046$ |
| | scConcept | 0.845 | 0.644 | $0.890 \pm 0.043$ |
| | scFoundation | 0.810 | 0.655 | $0.870 \pm 0.053$ |
| | scGPT | 0.773 | 0.500 | $0.707 \pm 0.061$ |
| | Geneformer-v2 | 0.685 | 0.363 | $0.687 \pm 0.061$ |
| Pancreas | scRep-30M | 0.794 | 0.687 | $0.890 \pm 0.021$ |
| | scRep-3M | 0.733 | 0.606 | $0.867 \pm 0.019$ |
| | STATE (SE-600M) | 0.813 | 0.655 | $0.860 \pm 0.027$ |
| | Stack | 0.835 | 0.719 | $0.920 \pm 0.015$ |
| | scConcept | 0.690 | 0.453 | $0.839 \pm 0.004$ |
| | scFoundation | 0.615 | 0.496 | $0.854 \pm 0.006$ |
| | scGPT | 0.657 | 0.442 | $0.681 \pm 0.025$ |
| | Geneformer-v2 | 0.363 | 0.156 | $0.634 \pm 0.014$ |
| Heart | scRep-30M | 0.829 | 0.844 | $0.818 \pm 0.008$ |
| | scRep-3M | 0.830 | 0.848 | $0.824 \pm 0.007$ |
| | STATE (SE-600M) | 0.831 | 0.847 | $0.790 \pm 0.020$ |
| | Stack | 0.807 | 0.807 | $0.818 \pm 0.008$ |
| | scConcept | 0.830 | 0.844 | $0.831 \pm 0.007$ |
| | scFoundation | 0.759 | 0.708 | $0.826 \pm 0.007$ |
| | scGPT | 0.736 | 0.662 | $0.736 \pm 0.017$ |
| | Geneformer-v2 | 0.677 | 0.547 | $0.747 \pm 0.028$ |

**Table S3:**
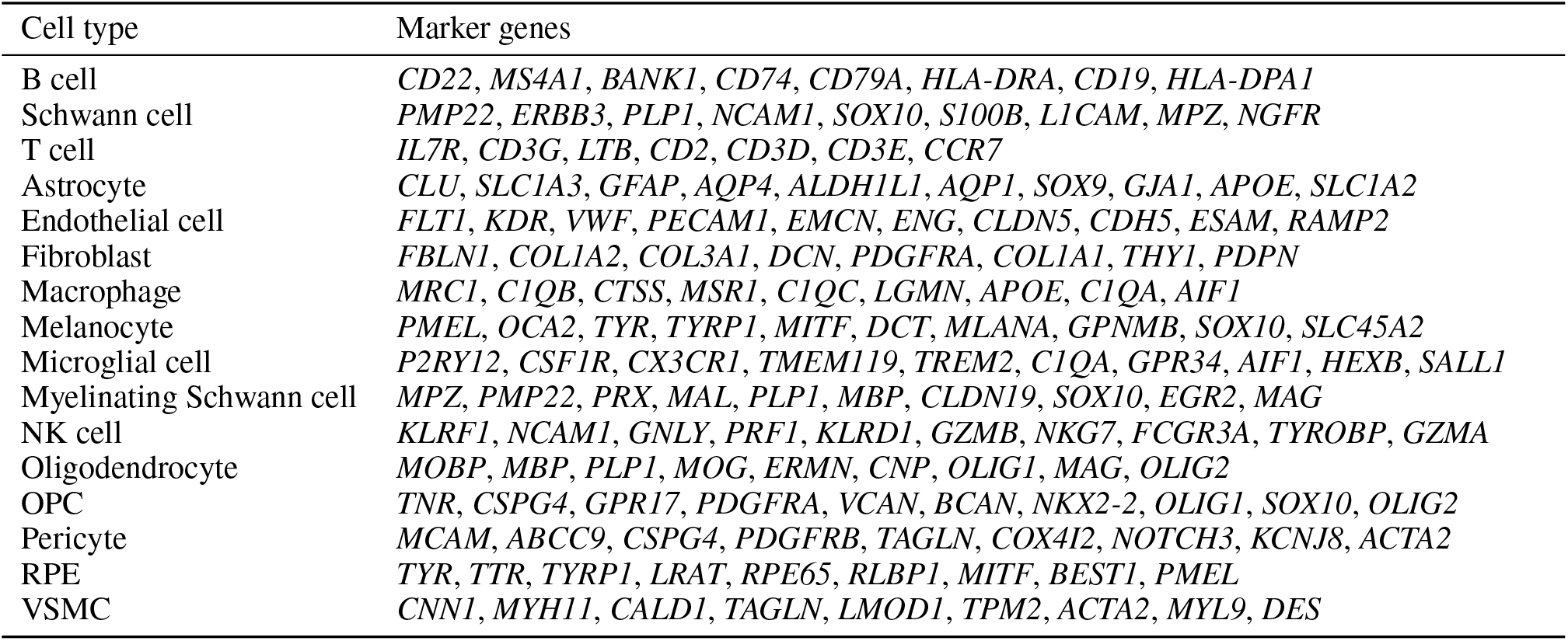
Cell-type marker genes used for the gene-importance analysis in the Eye dataset. Abbreviations: OPC, oligodendrocyte precursor cell; VSMC, vascular-associated smooth muscle cell; RPE, pigmented epithelial cell.

## References

[1] Abhinav K Adduri, Dhruv Gautam, Beatrice Bevilacqua, Alishba Imran, Rohan Shah, Mohsen Naghipourfar, Noam Teyssier, Rajesh Ilango, Sanjay Nagaraj, Mingze Dong, et al. Predicting cellular responses to perturbation across diverse contexts with state. BioRxiv, pages 2025–06, 2025.

[2] Mojtaba Bahrami, Alejandro Tejada-Lapuerta, Sören Becker, Fatemeh S Hashemi G, and Fabian J Theis. scconcept: Contrastive pretraining for technology-agnostic single-cell representations beyond reconstruction. bioRxiv, pages 2025–10, 2025.

[3] Rishi Bommasani, Drew A Hudson, Ehsan Adeli, Russ Altman, Simran Arora, Sydney von Arx, Michael S Bernstein, Jeannette Bohg, Antoine Bosselut, Emma Brunskill, et al. On the opportunities and risks of foundation models. arXiv preprint arXiv:2108.07258, 2021.

[4] Mathilde Caron, Hugo Touvron, Ishan Misra, Hervé Jégou, Julien Mairal, Piotr Bojanowski, and Armand Joulin. Emerging properties in self-supervised vision transformers. In 2021 IEEE/CVF international conference on computer vision (ICCV), pages 9630–9640. IEEE, 2021.

[5] Han Chen, Madhavan S Venkatesh, Javier Gómez Ortega, Siddharth V Mahesh, Tarak N Nandi, Ravi K Madduri, Karin Pelka, and Christina V Theodoris. Scaling and quantization of large-scale foundation model enables resource-efficient predictions in network biology. Nature Computational Science, pages 1–14, 2026.

[6] Ting Chen, Simon Kornblith, Mohammad Norouzi, and Geoffrey Hinton. A simple framework for contrastive learning of visual representations. In International conference on machine learning, pages 1597–1607. PmLR, 2020.

[7] The Tabula Sapiens Consortium*, Robert C Jones, Jim Karkanias, Mark A Krasnow, Angela Oliveira Pisco, Stephen R Quake, Julia Salzman, Nir Yosef, Bryan Bulthaup, Phillip Brown, et al. The tabula sapiens: A multiple-organ, single-cell transcriptomic atlas of humans. Science, 376(6594):eabl4896, 2022.

[8] Haotian Cui, Chloe Wang, Hassaan Maan, Kuan Pang, Fengning Luo, Nan Duan, and Bo Wang. scgpt: toward building a foundation model for single-cell multi-omics using generative ai. Nature methods, 21(8):1470–1480, 2024.

[9] Xiaoteng Cui, Qixue Wang, Junhu Zhou, Yunfei Wang, Can Xu, Fei Tong, Hongjun Wang, and Chunsheng Kang. Single-cell transcriptomics of glioblastoma reveals a unique tumor microenvironment and potential immunotherapeutic target against tumor-associated macrophage. Frontiers in Oncology, 11:710695, 2021.

[10] Alan DenAdel, Madeline Hughes, Akshaya Thoutam, Anay Gupta, Andrew W Navia, Nicolo Fusi, Srivatsan Raghavan, Peter S Winter, Ava P Amini, and Lorin Crawford. Evaluating the role of pretraining dataset size and diversity on single-cell foundation model performance. Nature Methods, 23(7):1447–1457, 2026.

[11] Jacob Devlin, Ming-Wei Chang, Kenton Lee, and Kristina Toutanova. Bert: Pre-training of deep bidirectional transformers for language understanding. In Proceedings of the 2019 conference of the North American chapter of the association for computational linguistics: human language technologies, volume 1 (long and short papers), pages 4171–4186, 2019.

[12] Mingze Dong, Abhinav Adduri, Dhruv Gautam, Christopher Carpenter, Rohan Shah, Chiara Ricci-Tam, Yuval Kluger, Dave P Burke, and Yusuf H Roohani. Stack: In-context learning of single-cell biology. bioRxiv, 2026.

[13] Jean-Bastien Grill, Florian Strub, Florent Altché, Corentin Tallec, Pierre Richemond, Elena Buchatskaya, Carl Doersch, Bernardo Avila Pires, Zhaohan Guo, Mohammad Gheshlaghi Azar, et al. Bootstrap your own latent-a new approach to self-supervised learning. Advances in neural information processing systems, 33:21271–21284, 2020.

[14] Laleh Haghverdi, Maren Büttner, F Alexander Wolf, Florian Buettner, and Fabian J Theis. Diffusion pseudotime robustly reconstructs lineage branching. Nature methods, 13(10):845–848, 2016.

[15] Minsheng Hao, Jing Gong, Xin Zeng, Chiming Liu, Yucheng Guo, Xingyi Cheng, Taifeng Wang, Jianzhu Ma, Xuegong Zhang, and Le Song. Large-scale foundation model on single-cell transcriptomics. Nature methods, 21 (8):1481–1491, 2024.

[16] Stephanie C Hicks, F William Townes, Mingxiang Teng, and Rafael A Irizarry. Missing data and technical variability in single-cell rna-sequencing experiments. Biostatistics, 19(4):562–578, 2018.

[17] Luni Hu, Hua Qin, Yilin Zhang, Yi Lu, Ping Qiu, Zhihan Guo, Lei Cao, Wenjian Jiang, Yixin Shen, Qianqian Chen, et al. Regformer: a single-cell foundation model powered by gene regulatory hierarchies. Nature Communications, 2026.

[18] Johannes Jakubik, Michael Vössing Niklas Kühl, Jannis Walk, and Gerhard Satzger. Data-centric artificial intelligence: J. jakubik et al. Business & Information Systems Engineering, 66(4):507–515, 2024.

[19] Jared Kaplan, Sam McCandlish, Tom Henighan, Tom B Brown, Benjamin Chess, Rewon Child, Scott Gray, Alec Radford, Jeffrey Wu, and Dario Amodei. Scaling laws for neural language models. arXiv preprint arXiv:2001.08361, 2020.

[20] Kasia Z Kedzierska, Lorin Crawford, Ava P Amini, and Alex X Lu. Zero-shot evaluation reveals limitations of single-cell foundation models. Genome Biology, 26(1):101, 2025.

[21] Jin Li, Jun Wang, Ignacio L Ibarra, Xuesen Cheng, Malte D Luecken, Jiaxiong Lu, Aboozar Monavarfeshani, Wenjun Yan, Yiqiao Zheng, Zhen Zuo, et al. Single-cell atlas of the transcriptome and chromatin accessibility in the human retina. Nature genetics, 58(2):418–433, 2026.

[22] Romain Lopez, Jeffrey Regier, Michael B Cole, Michael I Jordan, and Nir Yosef. Deep generative modeling for single-cell transcriptomics. Nature methods, 15(12):1053–1058, 2018.

[23] Mohammad Lotfollahi, F Alexander Wolf, and Fabian J Theis. scgen predicts single-cell perturbation responses. Nature methods, 16(8):715–721, 2019.

[24] Malte D Luecken, Maren Bü ttner, Kridsadakorn Chaichoompu, Anna Danese, Marta Interlandi, Michaela F Müller, Daniel C Strobl, Luke Zappia, Martin Dugas, Maria Colomé-Tatché, et al. Benchmarking atlas-level data integration in single-cell genomics. Nature methods, 19(1):41–50, 2022.

[25] Samuel W Lukowski, Camden Y Lo, Alexei A Sharov, Quan Nguyen, Lyujie Fang, Sandy SC Hung, Ling Zhu, Ting Zhang, Ulrike Grünert, Tu Nguyen, et al. A single-cell transcriptome atlas of the adult human retina. The EMBO journal, 38(18):EMBJ2018100811, 2019.

[26] Leland McInnes, John Healy, and James Melville. Umap: Uniform manifold approximation and projection for dimension reduction. arXiv preprint arXiv:1802.03426, 2018.

[27] Madhvi Menon, Shahin Mohammadi, Jose Davila-Velderrain, Brittany A Goods, Tanina D Cadwell, Yu Xing, Anat Stemmer-Rachamimov, Alex K Shalek, John Christopher Love, Manolis Kellis, et al. Single-cell transcriptomic atlas of the human retina identifies cell types associated with age-related macular degeneration. Nature communications, 10(1):4902, 2019.

[28] Aboozar Monavarfeshani, Wenjun Yan, Christian Pappas, Kenechukwu A Odenigbo, Zhigang He, Ayellet V Segrè, Tavé van Zyl, Gregory S Hageman, and Joshua R Sanes. Transcriptomic analysis of the ocular posterior segment completes a cell atlas of the human eye. Proceedings of the National Academy of Sciences, 120(34):e2306153120, 2023.

[29] Maxime Oquab, Timothée Darcet, Théo Moutakanni, Huy Vo, Marc Szafraniec, Vasil Khalidov, Pierre Fernandez, Daniel Haziza, Francisco Massa, Alaaeldin El-Nouby, et al. Dinov2: Learning robust visual features without supervision. arXiv preprint arXiv:2304.07193, 2023.

[30] CZI Cell Science Program, Shibla Abdulla, Brian Aevermann, Pedro Assis, Seve Badajoz, Sidney M Bell, Emanuele Bezzi, Batuhan Cakir, Jim Chaffer, Signe Chambers, et al. Cz cellxgene discover: a single-cell data platform for scalable exploration, analysis and modeling of aggregated data. Nucleic acids research, 53(D1): D886–D900, 2025.

[31] Alec Radford, Karthik Narasimhan, Tim Salimans, Ilya Sutskever, et al. Improving language understanding by generative pre-training. 2018.

[32] Rohit Rao, Rong Han, Sean Ogurek, Chengbin Xue, Lai Man Wu, Liguo Zhang, Li Zhang, Jian Hu, Timothy N Phoenix, Stephen N Waggoner, et al. Glioblastoma genetic drivers dictate the function of tumor-associated macrophages/microglia and responses to csf1r inhibition. Neuro-oncology, 24(4):584–597, 2022.

[33] Yanay Rosen, Yusuf Roohani, Ayush Agrawal, Leon Samotorčan, Tabula Sapiens Consortium, Stephen R Quake, and Jure Leskovec. Universal cell embeddings: A foundation model for cell biology. BioRxiv, pages 2023–11, 2023.

[34] Jun-ichi Satoh, Naohiro Asahina, Shouta Kitano, and Yoshihiro Kino. A comprehensive profile of chip-seq-based pu. 1/spi1 target genes in microglia. Gene regulation and systems biology, 8:GRSB–S19711, 2014.

[35] Oliver Stegle, Sarah A Teichmann, and John C Marioni. Computational and analytical challenges in single-cell transcriptomics. Nature Reviews Genetics, 16(3):133–145, 2015.

[36] Milena Stevanovic, Danijela Drakulic, Andrijana Lazic, Danijela Stanisavljevic Ninkovic, Marija Schwirtlich, and Marija Mojsin. Sox transcription factors as important regulators of neuronal and glial differentiation during nervous system development and adult neurogenesis. Frontiers in molecular neuroscience, 14:654031, 2021.

[37] C Claus Stolt, Anita Schlierf, Petra Lommes, Simone Hillgärtner, Torsten Werner, Thomas Kosian, Elisabeth Sock, Nicoletta Kessaris, William D Richardson, Veronique Lefebvre, et al. Soxd proteins influence multiple stages of oligodendrocyte development and modulate soxe protein function. Developmental cell, 11(5):697–709, 2006.

[38] Alejandro Tejada-Lapuerta, Anna C Schaar, Robert Gutgesell, Giovanni Palla, Lennard Halle, Mariia Minaeva, Larsen Vornholz, Leander Dony, Francesca Drummer, Till Richter, et al. Nicheformer: a foundation model for single-cell and spatial omics. Nature methods, 22(12):2525–2538, 2025.

[39] Christina V Theodoris, Ling Xiao, Anant Chopra, Mark D Chaffin, Zeina R Al Sayed, Matthew C Hill, Helene Mantineo, Elizabeth M Brydon, Zexian Zeng, X Shirley Liu, et al. Transfer learning enables predictions in network biology. Nature, 618(7965):616–624, 2023.

[40] Vincent A Traag, Ludo Waltman, and Nees Jan Van Eck. From louvain to leiden: guaranteeing well-connected communities. Scientific reports, 9(1):5233, 2019.

[41] Ashish Vaswani, Noam Shazeer, Niki Parmar, Jakob Uszkoreit, Llion Jones, Aidan N Gomez, Lukasz Kaiser, and Illia Polosukhin. Attention is all you need. Advances in neural information processing systems, 30, 2017.

[42] Hongzhi Wen, Wenzhuo Tang, Xinnan Dai, Jiayuan Ding, Wei Jin, Yuying Xie, and Jiliang Tang. Cellplm: Pretraining of cell language model beyond single cells. In International Conference on Learning Representations, volume 2024, pages 5649–5673, 2024.

[43] Yan Wu and Kun Zhang. Tools for the analysis of high-dimensional single-cell rna sequencing data. Nature Reviews Nephrology, 16(7):408–421, 2020.

[44] Fan Yang, Wenchuan Wang, Fang Wang, Yuan Fang, Duyu Tang, Junzhou Huang, Hui Lu, and Jianhua Yao. scbert as a large-scale pretrained deep language model for cell type annotation of single-cell rna-seq data. Nature machine intelligence, 4(10):852–866, 2022.

[45] Xiaodong Yang, Guole Liu, Guihai Feng, Dechao Bu, Pengfei Wang, Jie Jiang, Shubai Chen, Qinmeng Yang, Hefan Miao, Yiyang Zhang, et al. Genecompass: deciphering universal gene regulatory mechanisms with a knowledge-informed cross-species foundation model. Cell Research, 34(12):830–845, 2024.

[46] Daochen Zha, Zaid Pervaiz Bhat, Kwei-Herng Lai, Fan Yang, Zhimeng Jiang, Shaochen Zhong, and Xia Hu. Data-centric artificial intelligence: A survey. ACM Computing Surveys, 57(5):1–42, 2025.

